# A novel class of ultra-stable endospore appendages decorated with collagen-like tip fibrillae

**DOI:** 10.1101/2023.10.23.563578

**Authors:** Mike Sleutel, Ephrem Debebe Zegeye, Ann Katrin Llarena, Brajabandhu Pradhan, Marcus Fislage, Kristin O’Sullivan, Marina Aspholm, Han Remaut

## Abstract

Bacterial endospores are remarkable examples of biological resilience, representing a dormant and heavily fortified differentiation form capable of withstanding physical and chemical stressors detrimental to vegetative cells. In pathogenic firmicutes, spores also form an infectious particle and can take up a central role in the environmental persistence and dissemination of disease. A poorly understood aspect of spore-mediated infection is the fibrous structures or ‘endospore appendages’ (ENAs) that have been seen to decorate the spores of pathogenic Bacilli and Clostridia. New methodological approaches are opening an unprecedented window on these long enigmatic structures. Using cryoID, Alphafold modelling and genetic approaches we identify a novel class of ultra-robust ENAs formed by *Bacillus paranthracis*. We demonstrate that L-ENA are encoded by a three-gene cluster (*ena3*) that contains all components for the self-assembly of ladder-like protein nanofibers of stacked heptameric rings, their anchoring to the exosporium, and their termination in a trimeric ‘ruffle’ made of a collagen-like BclA paralogue. Phylogenomic analyses shows the *ena3* gene cluster as a mobile element with a polyphyletic distribution across pathogenic Bacilli.

## Introduction

Endospores formed by bacteria in the Firmicutes phylum are among the most durable lifeforms found in nature. Testament to their resilience is the spore-based panspermia research that has shown that spores would be able to withstand all the steps involved to traverse the solar system, including exposure to (i) g-forces associated with the ejection from and re-entry into earth’s orbit, (ii) high doses of radiation, and (iii) a combination of ultra-low vacuum and temperatures close to absolute zero. This cryptobiotic life form can bridge long periods that are not conducive to survival of the vegetative state, waiting for environmental stimuli to trigger germination and terminate the state of hibernation. This remarkable resilience can in part be traced back to the extraordinary robust layers that encase the genomic DNA, while maintaining the permeability for germinants that signal that the conditions are again conducive for metabolically active life (*1, 2*). For the endospore forming bacteria of the genera *Bacillus* and *Clostridium*, the genome is housed within a partially dehydrated central core that is encapsulated by a thick peptidoglycan layer, referred to as the cortex. The cortex, in turn, is covered by four consecutive protein layers that make up the spore coat: i.e., the basement layer, the inner and outer coat and the crust, comprising about 70 different proteins (*3, 4*). Basic models for the architectural layout of this molecular fortress have been proposed for *Bacillus subtilis* based on *in situ* localization experiments and genetic interaction networks, but a molecular understanding of this structure, or how it disintegrates during germination are largely missing.

Many species have an additional outermost layer called the exosporium. The exosporium is a flexible, glycosylated, para-crystalline sac-like structure that surrounds the spore, defining an interstitial volume between the crust and the exosporium. As the outermost spore layer, the exosporium mediates endosporés connection to the environment and the infected host (*5*). Interestingly, spores can be decorated with micrometer long filamentous structures called endospore appendages (ENA). First reported and classified for *Clostridium* spores in the late 1960s (*7–9*), followed by *Bacillus cereus* in the 1970s (*10*), ENAs can vary greatly in length, diameter and overall shape. They are present on spores of notable human and animal pathogens such as *Bacillus anthracis*, *B. cereus* (*10*), *Clostridium botulinum* (*9*) and *C. sordellii* (*11*), and seem mostly absent or unknown from saprophytic species, including *B. subtilis*. Pilus-like structures are well known from vegetative cells (*12*). In Gram-positive bacteria, pili are predominantly formed by the sortase-mediated covalent coupling and cell-wall anchoring of secreted LPxTG motif containing subunits via isopeptide bonds (*13–15*). These pili are primarily found in pathogenic species (i.e. *Corynebacterium dyphtheriae*, *Staphylococcus spp.*, *Enterococcus spp.*, *Streptococcus spp.* and *Bacillus* spp.), where they serve an adhesive function, binding to host tissues or mediating self-contact in multicellular communities. Endospores represent an important dissemination form in pathogenic *Bacillaceae*, so that similar to pili on vegetative cells, ENAs may mediate cell-cell and cell-tissue interaction. However, the resilience of ENAs to common proteomic approaches have hampered their molecular identification, in term preventing the genetic study of their function.

Due to their extreme physico-chemical robustness, ENAs have for decades resisted traditional proteomic identification methods (*16*) (N-terminal sequencing; peptide fingerprinting). This stalemate was recently broken by the virtue of cryoID, the direct molecular identification of proteinaceous structures through 3D reconstruction by cryo-electron transmission microscopy (cryoEM) (*17, 18*). Spores of the food poisoning outbreak strain *Bacillus paranthracis* NVH 0075-95 were found to display two structurally distinct ENAs, dubbed staggered and ladder-like or S- and L-ENA, respectively (*17*) (*18*). CryoID studies of S-ENA revealed it represents a novel pilus superfamily predominantly found in the *B. cereus* sensu lato group *(B. cereus* s.l.). S-ENA fibers are composed of two major subunits, Ena1A and Ena1B. Both proteins consist of a single DUF3992 domain comprising a β-jellyroll fold with an N-terminal extension. Ena1A and 1B subunits self-organize into micrometers long helical ultra-structures by lateral β-sheet augmentation, fortified by lateral and longitudinal inter-molecular disulfide bridges. Judging from negative stain transmission electron microscopy (nsTEM) images, S-ENA fibers seem to penetrate the exosporial sac and connect to the outer spore coat. Although the exact anchoring mechanism is unknown, we hypothesize that a third protein from the local *ena* gene cluster (Ena1C) could be involved in connecting S-ENA to the spore coat. Strikingly, S-ENAs terminate into 4 to 6 flexible, 2nm diameter, tip-fibrillae (hereafter referred to as *ruffles*) for which a genetic identification or structural elucidation remains absent. Phylogenomic analysis showed that *ena1/2* clusters are ubiquitous in pathogenic Bacilli (>90% occurrence in *B. cereus* s.l.) suggesting that S-ENAs could be hitherto overlooked virulence factors. Biomechanical studies suggest S-ENAs are implicated in self-adherence and the clustering of spores, albeit by an unknown mechanism (*19*).

Interestingly, the spores of NVH 0075-95 are decorated with a second type of shorter (typically < 1µm), 7nm diameter ENAs, that are tethered to the exosporium layer. Based on the typical ‘ladder-like’ patterning of these fibers in nsTEM images, we dubbed this sub-type of appendages ‘L-ENA’. In parallel with S-ENAs, L-ENA fibers are decorated with a single ruffle at their distal terminus. Currently, both the composition and ultrastructure of the L-ENA / ruffle complex are unknown. In this contribution, we use a combined cryoEM - genome mining approach to identify the major structural subunit of L-ENA (hereafter referred to as Ena3A) and solve the helical fiber ultrastructure to a resolution of 3.3 Å. Phylogenomic analysis indicates that *ena3A* is less prevalent than *ena1/2* gene clusters and is found predominantly on mobile genetic elements (i.e., on large plasmids and flanked by putative transposons). We show that *ena3A* is embedded in a three-gene cluster that holds all components for L-ENA assembly and anchoring to the exosporium. Based on phenotypic screening of knockouts strains and AlphaFold modelling, we identify the remaining two genes of the L-ENA gene cluster to encode for (i) an anchoring protein that tethers L-ENA to the exosporium and (ii) a collagen-like BclA-homologue that constitutes the ruffle protein.

## Results

### L-ENA fibers constitute a novel sub-class of disulfide cross-linked endospore appendages

The endospores of the food-poisoning outbreak strain *B. paranthracis* NVH 0075-95 are decorated with two types of ENA fibers, i.e., S-ENA and L-ENA (Fig.1a-b). Close inspection of the exosporium layer reveals that L-ENA fibers protrude from the brush-like hairy nap layer (Fig.1b) seemingly tethered to the exosporium. L-ENA fibers have an apparent diameter of 8nm and exhibit a ladder-like uranyl staining pattern in nsTEM with a typical recurring distance of 4.6nm between consecutive ladder segments. L-ENAs consistently terminate in a single 2nm diameter tip-fibrillum or ruffle, that consists of a stalk region (45nm in length) and a terminal knob domain (Fig.1c). In preparation of cryoEM data collection, we produced a sample enriched in ENA via shear-induced dislodging from the spore surface followed by a series of purification steps as described in (*17*). Next, a 1568 movie cryoEM dataset (60 frames, 0.766Å/pixel) was collected on a JEOL CRYO ARM 300 microscope, yielding a 5.8Å resolution (FSC=0.143 criterion) reconstructed cryoEM-volume of L-ENA fibers with C7 symmetry (see Fig.1e and Supporting Figure 1). The refined helical parameters are a twist of 17.04°, and a rise of 43.82Å. We attribute the limited resolution of the final reconstruction to (i) the sparsity of images that contained L-ENA fibers (414 micrographs, i.e., 26% of the total dataset), and (ii) the innate flexibility of the L-ENA fibers (discussed below). The limited resolution precluded us to *de novo* build an atomic model of L-ENA but allowed us to identify the fold of the constituent monomers. Indeed, an exploratory rigid body docking exercise using Ena1B as template model (extracted from PDB: 7A02) taught us that the L-ENA subunits appeared structurally homologous to Ena1A/B and therefore likely represented members of the DUF3992 family. A careful search of the NVH 0075-95 genome (GCA_027945115.1) with HMMsearch 3.0 (*20*) using a hidden Markov model generated from a dataset of Ena1/2 protein sequences as query led to the identification of a fourth gene outside the *ena1a-c* cluster encoding for a DUF3992 domain-containing protein. As such, we reasoned that WP_017562367 represented the candidate L-ENA subunit and ordered its coding sequence as a synthetic gene, and cloned it into pET28a for cytoplasmic expression in *E. coli* C43 (DE3), similar to the approach taken for recombinant S-ENA production (*17*). To probe for the presence of putative L-ENA fibers, the insoluble fraction (after lysozyme treatment and 1% SDS extraction) of the *E. coli* lysate was analyzed using nsTEM. The obtained micrographs confirmed the presence of micron-sized, 7nm diameter fibers with a characteristic ladder-like pattern at 4.6nm intervals (Fig.1f). Notably, the recEna3A fibers did not have any ruffles on their termini. This result served as an initial confirmation that WP_017562367 is the major subunit of L-ENAs found on *B. paranthracis* NVH 0075-95. To reflect the fact that WP_017562367 is a member of a new *ena* gene cluster, we propose to name it *ena3a*, following the convention used for the *ena1a-c* (e.g., *B. cereus* type *S-ENA*) and *ena2a-c* (e.g., *B. thuringiensis* type *S-ENA*) (*17*).

**Figure 1:**
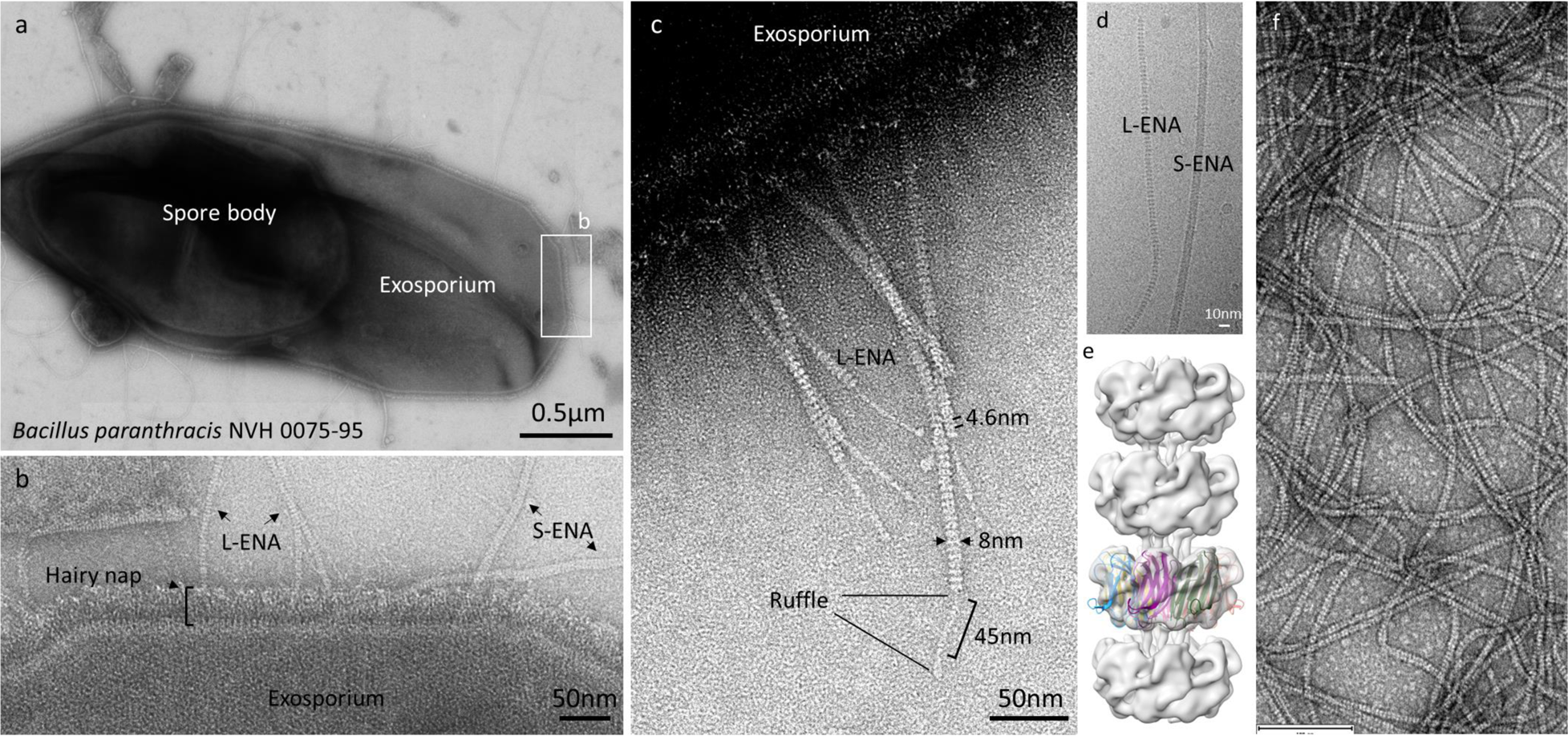
Negative stain transmission electron microscopy images of the endospores of the food-poisoning outbreak strain *B. paranthracis* NVH 0075-95: (a) Composite montage of a single endospore with a central spore body and the surrounding exosporial sack, (b) High-magnification image of the exosporium decorated with the hairy nap and two types of endospore appendages (L- and S-ENA), (c) L-ENA fibers anchored to the exosporium, exhibit a ‘ladder-like’ pattern (4.6nm intervals), are terminally decorated with single tip-fibrillae, i.e., ruffles. Ruffles consist of a 45nm long stalk that terminates into a globular head domain, (d) cryoEM image of isolated ENA fibers, (e) reconstructed cryoEM-volume of L-ENA fibers (5.8Å at FSC=0.143) with rigid body docking of 7 copies of Ena1B_19-117_, (f) nsTEM image of recEna3A produced in and purified from the cytoplasm of *E. coli*.

Given the relative ease of production and purification of recEna3A fibers in comparison to the L-ENA purification from the natural source, we proceeded to collect a 10886 movies cryoEM dataset of vitrified recEna3A fibers (Supporting Figure 2) leading to a 3.3 Å global resolution (FSC=0.143 criterion) cryoEM volume after helical refinement using C7 symmetry in CryoSPARC v4.0.3 (Fig.2). The refined helical parameters are a twist of 18.5°, and a rise of 44.97Å. The recEna3A cryoEM volume reveals an axial stacking of heptameric Ena3A rings that rotate 18.5° clockwise relative to each other (Fig.2a). Each ring is composed of seven Ena3A molecules that interface laterally via β-sheet augmentation. The rings encircle a central, hollow volume which we refer to as the ring lumen (Fig.2b). Consecutive rings interlock with each other using the N-terminal extensions of the upper ring docking into the lumen of the ring below (Fig.2c). We refer to the first 14 N-terminal residues as the N-terminal connector (i.e., Ntc) in analogy to the N-termini of Ena1A/B subunits that interlock from above with subunits i-9 and i-10 of the neighboring helical staircase (*17*). Note that the orientation of the L-ENA fiber in Fig. 2 is such that the top rings represent the distal, ruffled end of the fiber, and that the bottom rings form the pointy end of the fibers, proximal to the spore surface.

**Figure 2:**
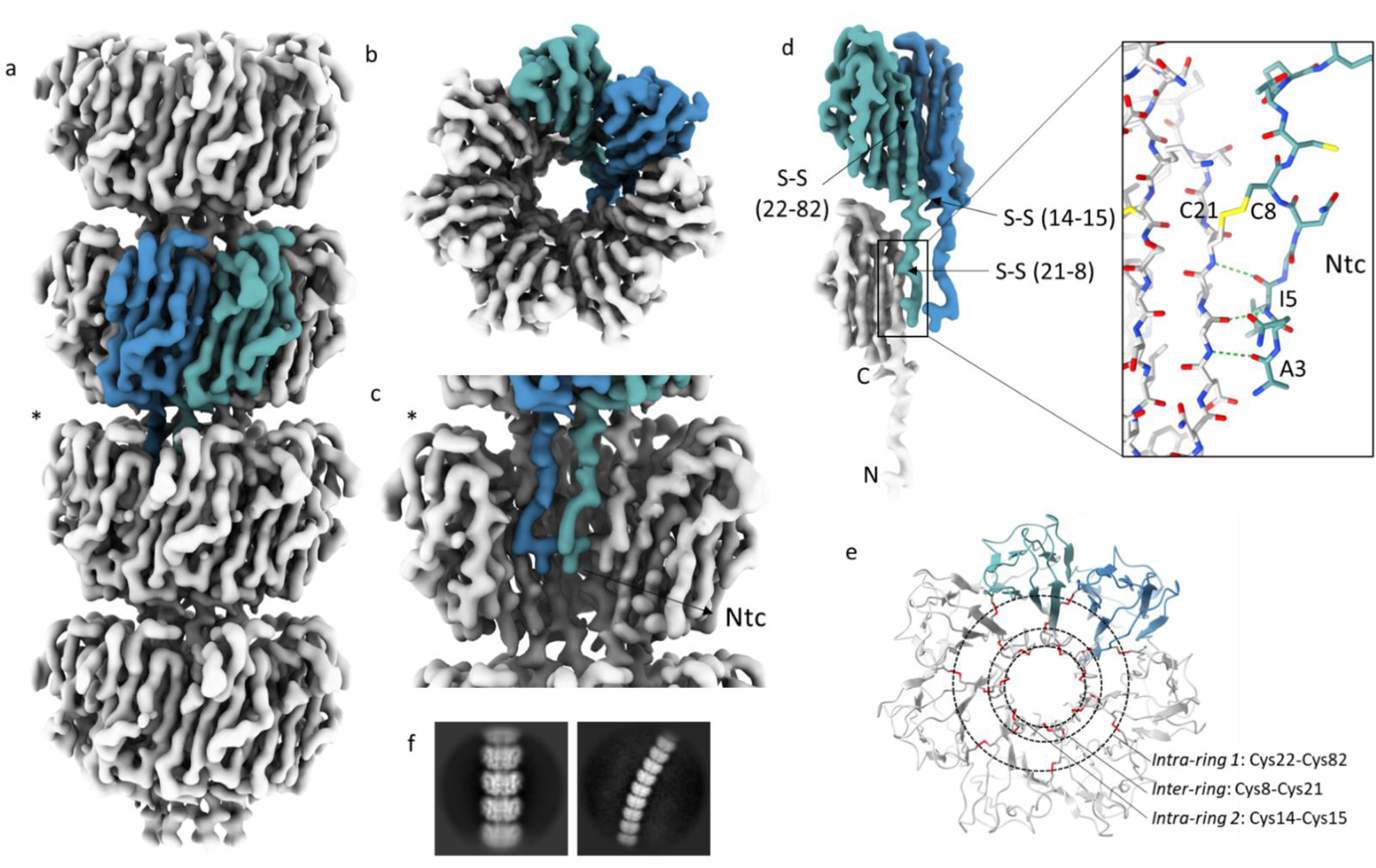
CryoEM volume of recEna3A L-ENA fibers: (a) Helical ultrastructure of L-ENA determined at 3.32 Å global resolution (cut-off criterion at FSC=0.143 Å^-1^) revealing an axial stacking of heptameric Ena3A rings that rotate 18.5° clockwise relative to each other. Two neighboring Ena3A monomers are colored in blue and cyan, (b) top-view of a single ring with a highlight of the β-sheet augmentation between the blue and cyan subunit, (c) zoom-in of a single ring - for clarity, two subunits were removed from the ring highlighted with an asterisk in (a) - showing the docking of the N-terminal connectors that covalently tether the ring docked above, (d) Highlight of the lateral intra-ring contacts: two types of disulfide bridges (Cys14-Cys15; Cys22-Cys82) exist within two neighboring subunits. In turn, each subunit connects to a subunit in the ring below via their Ntc (Cys8-Cys21; inset in stick representation), (e) On axis top-view of the L-ENA fiber model in cartoon with disulfide bridges shown in red; (f) 2D class averages of straight and curved L-ENA segments covering 4 and 9 rings, respectively,

Close inspection of the cryoEM map revealed covalent cross-links between neighboring Ena3A subunits at three distinct locations (Fig.2d). After manual building of the final L-ENA model, we identified these contacts to be 3 types of inter-molecular disulfide bridges. Two disulfide bridges (i.e., Cys22-Cys82 and Cys14-Cys15) serve to reinforce the lateral contacts within a single ring, whereas one disulfide bridge (i.e., Cys8-Cys21) confers longitudinal coupling between the lumen and the Ntcs of consecutive rings. Given the heptameric nature of the structure, each ring segment will contain 21 inter-molecular disulfide bridges that are organized along three concentric rings centered on the fiber axis (Fig.2e). Next we tested the stability of recombinant L-Ena fibers by subjecting the fibers to a series of physical (autoclaving) and/or chemical treatments (8M urea, SDS, proteinase K, formic acid). Following the respective treatments, we perform nsTEM imaging of the recovered fibers and determine the 2D class average to gauge the intactness of the fibers (Supporting Figure 4). Remarkably, the 2D class average images of all treatments (apart from the 100% formic acid) are very similar to the 2D class averages obtained from *ex* vivo purified fibers. We were not able to produce 2D class averages for the 100% formic acid (FA) treated sample, likely indicating that the Ena3A monomers had (partially) unfolded. Interestingly, the FA-treated fibers did not depolymerize, suggesting that disulphide bonds had not been reduced. We conclude that the extensive hydrogen bonding between the Ena3A subunits combined with the covalent cross-links underlies the physico-chemical robustness of the L-ENA fibers.

L-ENA marries this extreme stability with a remarkable flexibility as judged from the regions of high local curvature in the nsTEM micrographs and 2D class average images obtained during cryoEM processing (Fig.1f). To illustrate the point further, we manually selected and extracted curved L-ENA fiber segments from the motion-corrected micrographs and performed 2D classification (Supporting Figure 3a). To resolve the underlying structural heterogeneity, we performed a 3D variability analysis using CryoSPARC (Supporting Figure 3b). The resulting volume series (processed in ChimeraX 1.4 (*21, 22*) and exported as Supporting Movie 1) provides further molecular insights into the L-ENA flexibility. Ena3A rings exhibit a rocking motion normal to the fiber axis with the respective hinge points centered on the Ntcs. Hence, L-ENA flexibility can be traced back to (i) the spatial separation between consecutive rings thereby providing the possibility for local ring displacement without introducing steric clashes, (ii) combined with the intrinsic flexibility of the N-terminal connectors.

Ena3A subunits consist of a typical jellyroll fold (*23*) comprised of two juxtaposed β-sheets containing strands BIDG and CHEF (Fig.3). As mentioned earlier, the jellyroll domain is preceded by a flexible 14-residue Ntc that mediates inter-ring coupling. The backbone (420 atoms) root-mean squared displacement (RMSD) between Ena1B and Ena3A is 3.7Å even though the sequence identity is only 28.4% (Supporting Figure 5a-c). Despite the structural similarities at the fold level, S- and L-ENA have a markedly different quaternary architecture (Figure 3a-c). Ena1B subunits interact laterally via β-sheet augmentation (Supporting Figure 5d-e) with a non-zero (i.e., 3.2Å) vertical offset, making a 28° angle relative to the fiber axis. This in turn leads to a helical stacking of Ena1B monomers yielding an S-ENA fiber with a diameter of 110Å. Ena3A subunits form a similar dimer interface (see the residues marked with an asterisk in Supporting Figure 5b) that follows the same register between the interfacing strands G and C (Supporting Figure 5f-g). Contrary to Ena1B though, neighboring Ena3A subunits lie within the same plane (i.e., no vertical offset) thereby producing closed ring-like structures. We attribute this difference in axial displacement to the relative tilts that the subunits make with respect to the fiber axis, i.e., 28° and 14° for Ena1B and Ena3A, respectively (Fig. 3c and f). L-ENA fiber biogenesis therefore likely proceeds via docking and covalent locking (via the Cys8-Cys21 disulfide) of fully formed rings, whereas S-ENA fibers elongate via integration of successive monomers.

**Figure 3:**
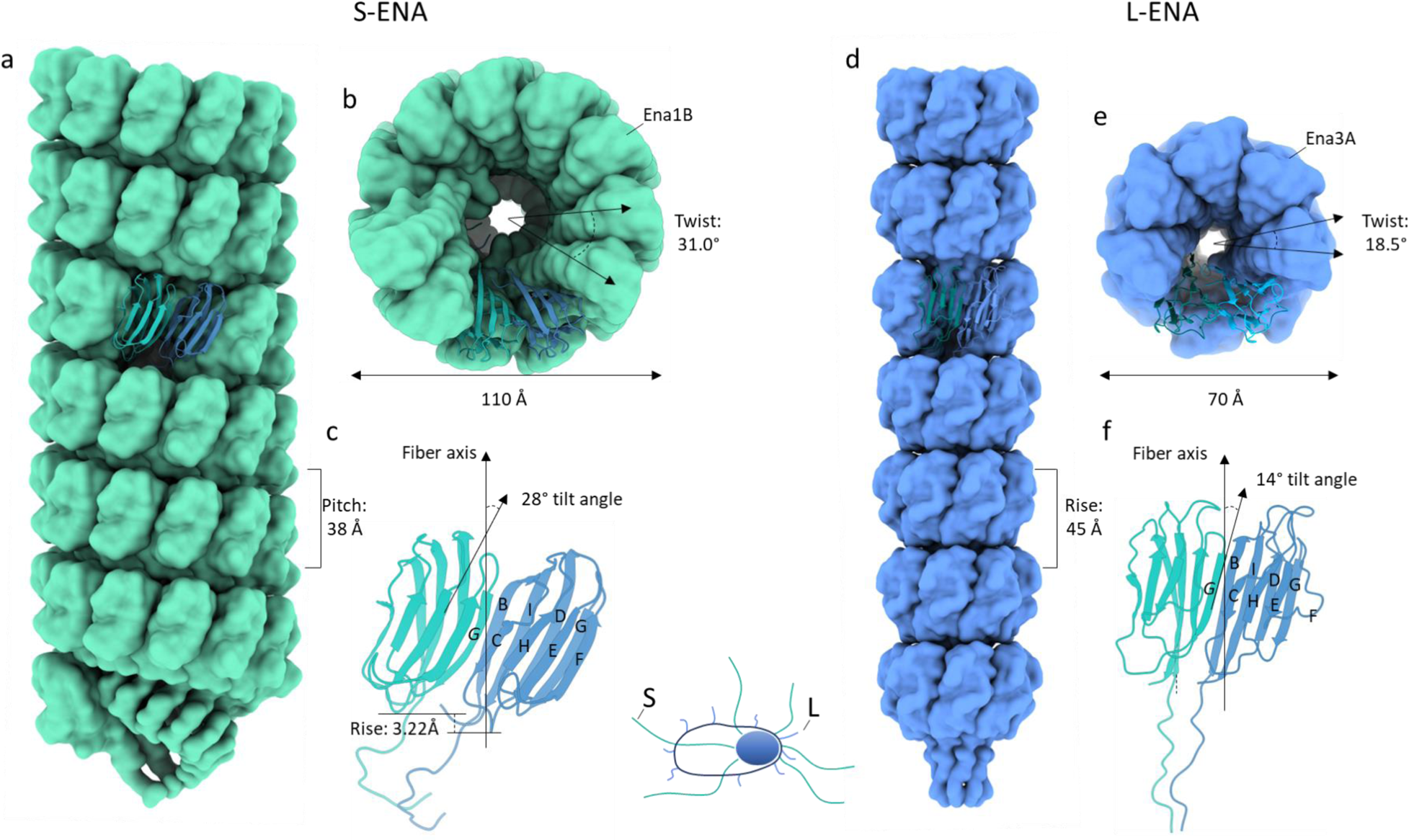
Structural comparison between the S- and L-ENA fiber architectures: (a) Side and (b) on-axis view of an S-ENA fiber (pdb-id: 7A02) composed of Ena1B subunits. Helical parameters: Rise: 3.22Å, Twist: 31.0°, Pitch: 38Å), (c) Dimeric contacts of Ena1B subunits via β-sheet augmentation at the interface between the G and C strands. Subunits are tilted 28° with respect to the fiber axis. This out-of-plane interaction leads to a helical stacking of Ena1B monomers, (d) Side and (e) on-axis view of an L-ENA fiber (pdb-id: 8C50) composed of Ena3A subunits. Helical parameters: Rise = pitch: 45Å, Twist: 18.5°, C7), (f) Dimeric contacts of Ena3A subunits via β-sheet augmentation at the interface between the G and C strands. Although subunits are tilted 14° with respect to the fiber axis, their contacts remain in-plane yielding a heptameric ring.

### The L-ENA subunit, exosporium anchoring protein and ruffle protein are encoded in a three-gene cluster

Inspection of the GCA_027945115 genome reveals that *ena3A* (PGS39_28750) is embedded in a three-gene cluster on the NZ_CP116205 plasmid, hereafter referred to as the *ena3* gene cluster (Figure 4). In this cluster, *ena3A* is preceded by the genes PGS39_28740 and PGS39_28745. For reasons discussed below, we suggest naming these two genes, i.e. *cotZ* and *l-bclA*, respectively. A primary sequence analysis of CotZ (WP_048548726.1) using Interpro (*24*) shows that it is composed of an N-terminal CotZ/Y domain and a C-terminal Ena core (DUF3992) domain (see the domain organization and corresponding AF2 prediction in Fig.4b). In turn, L-BclA (WP_271292911.1) consists of an N-terminal collagen-like region followed by a C-terminal domain that belongs to the *Bacillus* collagen-like protein of *anthracis* family, i.e. the head domain BclA-C of one of the main substituents of the hairy nap layer (*25*). In pursuit of the biological roles of *cotZ* and *l-bclA*, we made individual knockouts strains (Δ*cotZ* and Δ*l*-*bclA*) and investigated their respective endospores by nsTEM (Fig. 5). CotZ depleted spores are devoid of L-ENA fibers coupled to the exosporium but are otherwise morphologically identical to wild-type NVH 0075-95 spores. Careful inspection of the grid areas lead to the identification of detached L-ENAs in the spore supernatant, with ruffles present, suggesting that CotZ mediates the connection of L-ENA to the exosporium. Conversely, Δ*l-bclA* spores have L-ENAs present on the exosporium, but the fibers lack the distal ruffle. We do note that the ruffles remained present on the S-ENA fibers, suggesting that *l-bclA* specifically encodes for the L-ENA ruffle protein. As expected, *ena3a^-^* spore samples were completely devoid of L-ENA fibers, be it spore attached or detached. Complementation of the Δ*cotZ*, Δ*l-bclA* and Δ*ena3A* mutants with a low copy plasmid (pHT315) containing the respective genes, restored the corresponding spore phenotypes to that of the wild type strain (Fig. 5).

**Figure 4:**
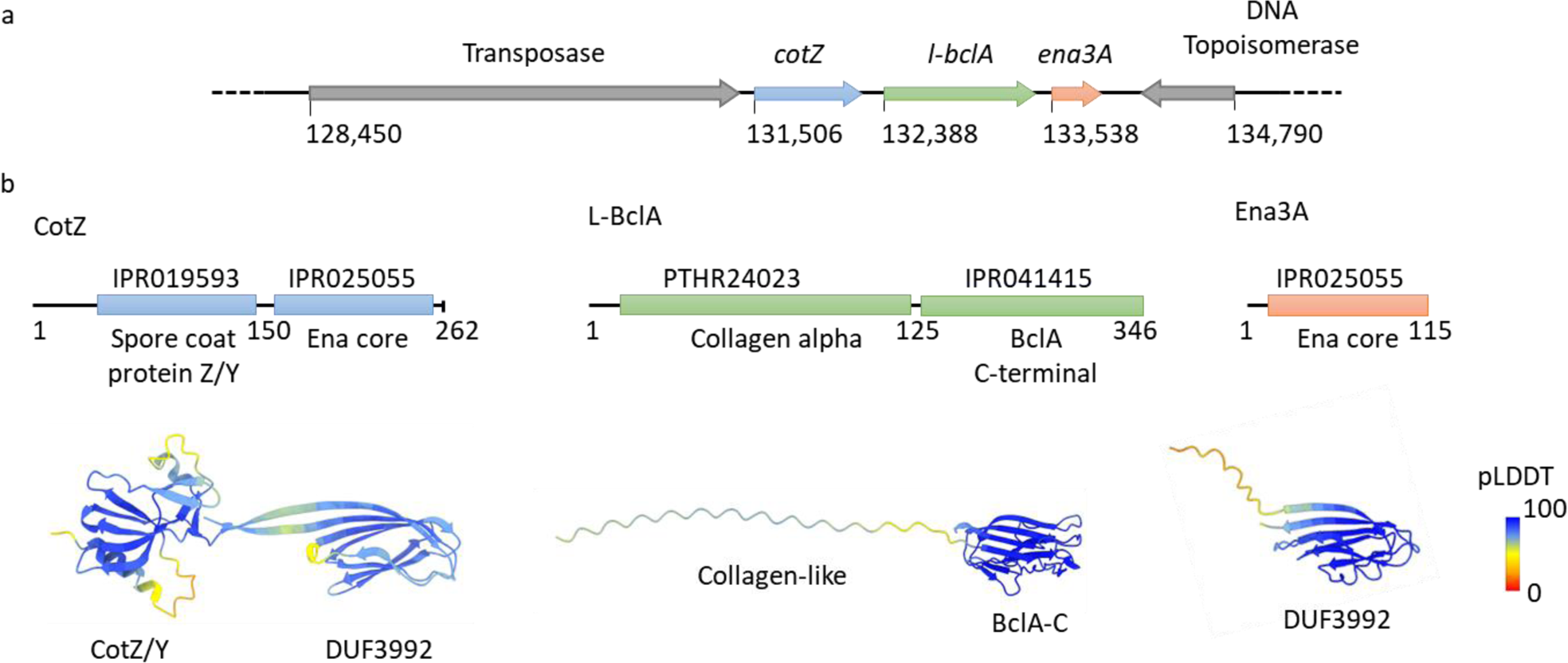
Genetic organization of the L-ENA gene cluster: the *ena3A* gene is embedded in a three gene cluster, preceded on the CP116205.1 plasmid by two genes here annotated as *cotZ* and *l-bclA*, (b) Domain organization of the L-ENA proteins as identified by Interpro (10.1093/nar/gkac993). CotZ: consists of an N-terminal spore coat protein Z/Y domain and a C-terminal Ena core domain (DUF3992), L-BclA: consists of an N-terminal collagen-like domain and a C-terminal BclA-C domain, Ena3A: consists of a single Ena core domain (DUF3992). AlphaFold2 predictions are shown in cartoon representation, colour coded according to the pLDDT score.

**Figure 5:**
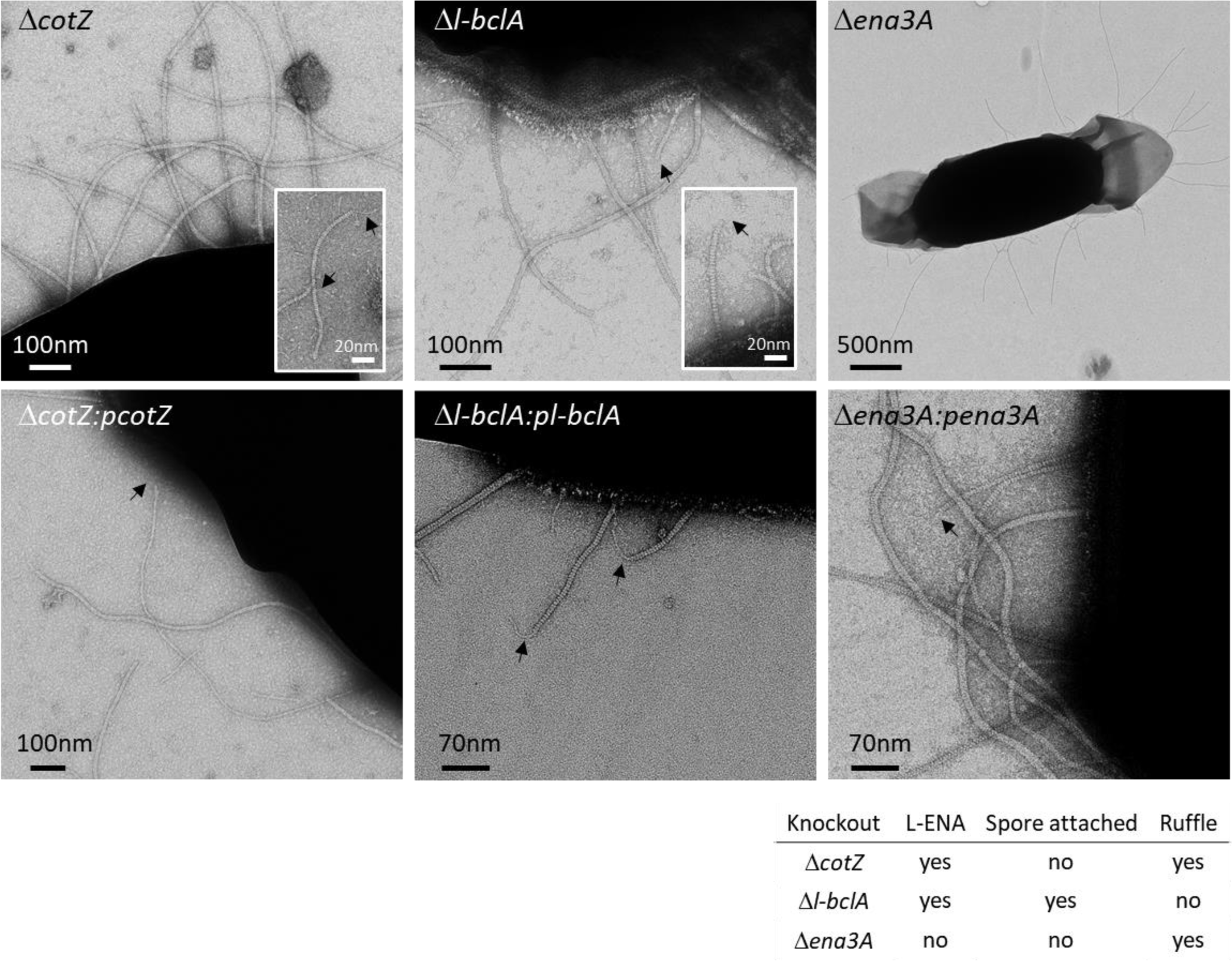
Phenotypic read-out of L-ENA mutants: nsTEM micrographs of the respective L-ENA operon deletion mutants.

### CotZ forms an exosporium-embedded anchoring proteins for L-ENA

To gain further insights into the molecular mechanisms of L-ENA anchoring and ruffle formation, we performed Alphafold2 (AF2) modelling (*26–28*). First, we tested for the plausibility of a Ena3A-CotZ complex formation. In Supporting Figure 6 we show the AF2 multimer v1.2 model of an Ena3A-CotZ dimer, which had an overall pLDDT score of 82.6 and a ptmscore of 0.73, which is in support of the hetero-dimer hypothesis. As expected, Ena3A interfaces with CotZ via its C-terminal Ena-core domain in a manner that mimics the Ena3A dimer interface found in the L-ENA structure (Supporting Figure 6e-f). This is somewhat surprising given the low sequence identity (17.4%) between the Ena3A and the CotZ Ena-core domain (i.e. residues 157-262). Inspection of the pairwise sequence alignment (Supporting Figure 6d) between Ena3A and CotZ learns that key Ena3A residues (C82, S84 and T86) involved in lateral subunit contacts are conserved in CotZ (C224, S226 and T228). In fact, AF2 predicts a disulfide bridge between CotZ cys224 and Ena3A cys22, which mimics the intra-ring L-ENA S-S bridge between cys22 and cys82 (Supporting Figure 6d; Fig. 2d-e). Despite the low sequence identity in Ena3A and the CotZ Ena-core, the Ena3A-Ena3A and Ena3A-CotZ contacts are highly equivalent (Supporting Figure 6d-g), in line with the general mechanism of β-sheet augmentation, which is primarily driven by shape complementarity and backbone H-bonding and is relatively insensitive to the amino acid sequence in the paring β-strands (*29*).

In addition, we looked at the AF2 prediction of a putative CotZ-ExsY dimer. ExsY (WCA20099.1) is one of the major constituents of the exosporium and shares structural similarities to the C-terminal CotZ/Y domain of the *cotZ* gene in the *ena3* gene cluster, albeit at low sequence identity, i.e., 30.8% (Supporting Figure 7 a,b,f). Our assumption based on the *cotZ^-^* phenotype and the Ena3A-CotZ AF2 model was that CotZ could also be a potential binding partner of ExsY and, in doing so, act as an exosprium-embedded anchor point for L-ENA fibers. Indeed, the AF2 multimer model of a CotZ-ExsY dimer (with a relatively high pLDDT score of 80.2 and a ptmscore of 0.71) shows an ExsY-CotZ coupling via the N-terminal CotZ_1-156_ domain with ExsY, lending further credence to the supposition that CotZ serves as a connective bridge between the paracrystalline exosporium and L-ENA. In analogy to the CotZ-Ena3A dimer, the predicted CotZ-ExsY contact mimics the homomeric ExsY contacts (Supporting Figure 7c,d,e,g) that we observe in a high confidence AF2 multimer prediction of an ExsY hexamer (pLDDT=81.3; ptmscore=0.86), i.e., the base building unit of the exosporium lattice.

### The L-ENA ruffle is formed by the collagen-like protein L-BclA

Next, we focused on L-BclA, a putative collagen-like protein that resembles BclA, the major glycoprotein of the hairy nap found to cover the exosporium of most *B. cereus s.l.* species (*25, 30*). In Supporting Figure 8a-b, we present the AF2 multimer model of an L-BclA_94-336_ homotrimer that consists of the C-terminal trimerization domain and a segment of the collagen-like triple helix. The corresponding pLDDT value of 94 and ptmscore of 0.9 support the accuracy of the predicted fold, as well as the supposed trimeric stochiometry in analogy to the crystal structure (PDB: 1WCK) of BclA-C of *B. anthracis*. Based on this model, we predict the lateral dimension of the C-terminal domain to be approximately 4nm, and for the triple collagen helix we project an average length of 2.8 Å per residue (see discussion below; Supporting Figure 8a). The all-atom RMSD between L-BclA-CTD and BclA-C is 3.55 Å despite the low sequence identity of 22.1%, demonstrative of significant structural homology (Supporting Figure 8c). Tan and Turnbough showed that the attachment of BclA to the basal layer protein BxpB of the exosporium is dependent on (i) proteolytic removal of the first 20 residues from the N-terminus and (ii) the presence of an N-terminal submotif -hereafter referred to as the exosporium leader sequence (interpro: IPR0212010; Supporting Figure 8d) - in front of the collagen-like region (*31*). Inspection of the N-terminal region of L-BclA demonstrates the absence of an exosporium leader sequence, which indicates that this collagen-like protein is likely not targeted to the exosporium.

A notable feature of collagen-like proteins is that the number of residues that comprise the collagen-stalk region in the primary sequence translates proportionally to the axial dimension of the folded, trimeric entity simply due to the extended nature of the collagen fold (*30*). To that end, we searched the GCA_027945115.1 genome for the ‘local’ orthologue of BclA (uniprot: Q81JD7) and compared its primary sequence to L-BclA (Supporting Figure 8e). BclA_NVH 0075-95_ (WCA20088.1) has a collagen-like region of 125 residues, whereas the corresponding domain of L-BclA spans 173 residues. Based on the calibration discussed in Supporting Figure 8a, this translates into a predicted fully extended length of a folded trimeric entity of 39 nm and 52 nm for BclA and L-BclA, respectively. These predictions are in agreement with experimental measurements (average height of the hairy nap: ∼36nm and average length of L-ENA ruffle: ∼50nm) from nsTEM micrographs (Supporting Figure 8f). To further understand the topology of the L-ENA/L-BclA complex, we performed a single round of 2D classification based on particles that were manually picked from the cryoEM dataset that was collected on *ex vivo* isolated ENA fibers, focusing on L-ENA termini that were decorated with ruffles (Supporting Figure 8g). This class average shows that the ruffle docks into the lumen of the Ena3A ring at the apex of the L-ENA fiber, i.e. mimicking the inter-ring docking mechanism that exists along the body of the fiber. We therefore compared the Ntc of Ena3A to the N-terminus of L-BclA (Supporting Figure 8d). Although no particular sequence homology could be detected, both sequences contain a double cysteine motif at positions 8 and 13, respectively. As cysteine 8 mediates coupling of the Ena3A Ntc to the ring lumen, we speculate that the L-BclA C_13_C_14_ motif could be involved in a similar coupling mechanism.

### Distribution and expression of *ena3A, cotZ* and *l-bclA*

The *ena3* gene cluster is rare: it was found in only 62 organisms through a remote search of the entire NCBI RefSeq non-redundant protein database using cblaster (CAGECAT v. 1.0). Assemblies of sufficient quality were downloaded and appended to a representative database of genomes of the *Bacillus* genus. The proportion of genomes carrying an *ena3* gene cluster was indeed low; only 9.5% (n=62/656) of the *B. cereus s. l.* genomes, of which 51 were *B. cereus* (n=51/126), four *B. thuringiensis* (n=4/52), one *B. anthracis* (n=1/63), one *B. paranthracis* (n=1/4), one *B. toyonensis* (n=1/204), two *B. mobilis* (n=2/5), and two *Bacillus* sp. (2/4) (Supporting Figure 9). Most genomes had one copy of the gene cluster (n=53), while nine strains had paralogs, carrying two (n=5, *B. cereus (3)*, *Bacillus* sp. *(2)*), three (n=2, *B. cereus*), four (n=1, *B. thuringiensis*) and five (n=1, a *B. thuringiensis*) copies. The *ena3* gene cluster was not found in *B. subtilis* or other saprophytic Bacilli (see Supporting Data).

Thirteen genomes with an L-ENA gene cluster were complete and closed, and the genomic location of the gene cluster could be inspected. The *ena3* gene cluster was located either on the chromosome (n=7 genomes), on plasmids (n=1 genome) or were found as paralogs on one or more plasmids and on the chromosome (n=5 genomes). Many of the isolates carrying the *ena3* gene cluster were from outbreaks of bloodborne infections in hospitals in Italy and Japan (*32, 33*).

The *ena3* gene cluster of NVH 0075-95 is flanked by an incomplete topoisomerase upstream and a complete Tn3 family transposase upstream. This gene synteny was found in only two other strains in the cblaster/clinker analysis: *B. cereus* AFS093282 (NZ_NVMQ01000017.1) and *B. pacificus* strain BC444B (accession NZ_JAOPRQ010000006.1). In other strains, the L-ENA gene cluster was flanked by frameshifted versions of Tn3 family transposases, tyrosine-type recombinase/integrase and site-specific integrases. A few strains had a shorter Tn3 family transposase upstream of the L-ENA gene clusters. Taken together, the L-ENA gene cluster is/or has been located on a transposon and is part of the mobilome of the *B. cereus s.l.* group, accounting for the polypheletic distribution of this gene cluster in the population.

To determine the expression of L-ENA genes (*cotZ*, *bclA* and *ena3A*), NVH0075-95 was cultured in sporulation medium for 16 hours, and cDNA prepared from culture samples collected at four-hour intervals after inoculation (4, 8, 12 and 16 hours) and analyzed by PCR. Using primers that specifically amplify open reading frames across *cotZ*→*l-bclA* and *l-bclA*→*ena3A* (Figure 6a), we detected a ∼587 bp PCR product across *l-bclA*→*ena3A*, but not for *cotZ*→*l-bclA* (Figure 6b). This suggests that *l-bclA* and *ena3A* are expressed bicistronically. Notably, no *l-bclA-ena3A* transcript was detected during the first 8 hrs of cultivation, which represents the vegetative growth phase (Fig. 6b). Consistent with the PCR result, the data from a qPCR analysis indicated that the L-ENA genes are overexpressed exclusively during the sporulation phase (12 and 16 hours) (Fig. 6c). Notably, a ∼13000, ∼1200 and ∼40-fold increase in the expression of *ena3A*, *l-bclA* and *cotZ*, respectively, was evident in the samples collected at 12 hours after inoculation.

**Figure 6:**
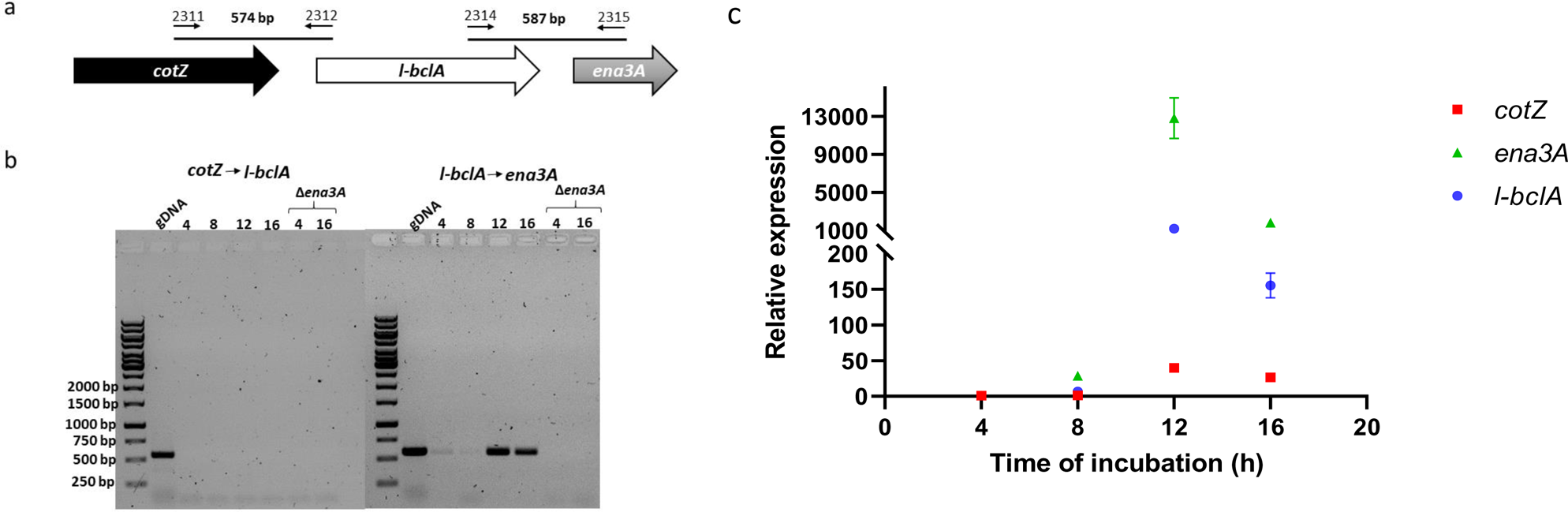
Expression of L-ENA genes is concomitant with sporulation. (a) Chromosomal organization of *cotZ*, *l-bclA* and *ena3A*, and primers used in standard PCR using cDNA template; (b) The L-ENA genes *l-bclA* and *ena3A* form an operon. Agarose gel electrophoresis (1%) of PCR products conducted on cDNA from NVH0075-95 culture (4, 8, 12 and 16 h). Primer combinations and expected product sizes are shown in (a). Genomic DNA (gDNA) from NVH0075-95 and cDNA prepared from its isogenic Δ*ena1ABC*/Δ*ena3A* mutant were used as positive and negative controls for the PCR, respectively; (c) Expression of L-ENA genes relative to *rpoB* during vegetative growth and sporulation. Error bars represent the standard deviation of three independent experiments, each with three technical replicates.

**Figure 7:**
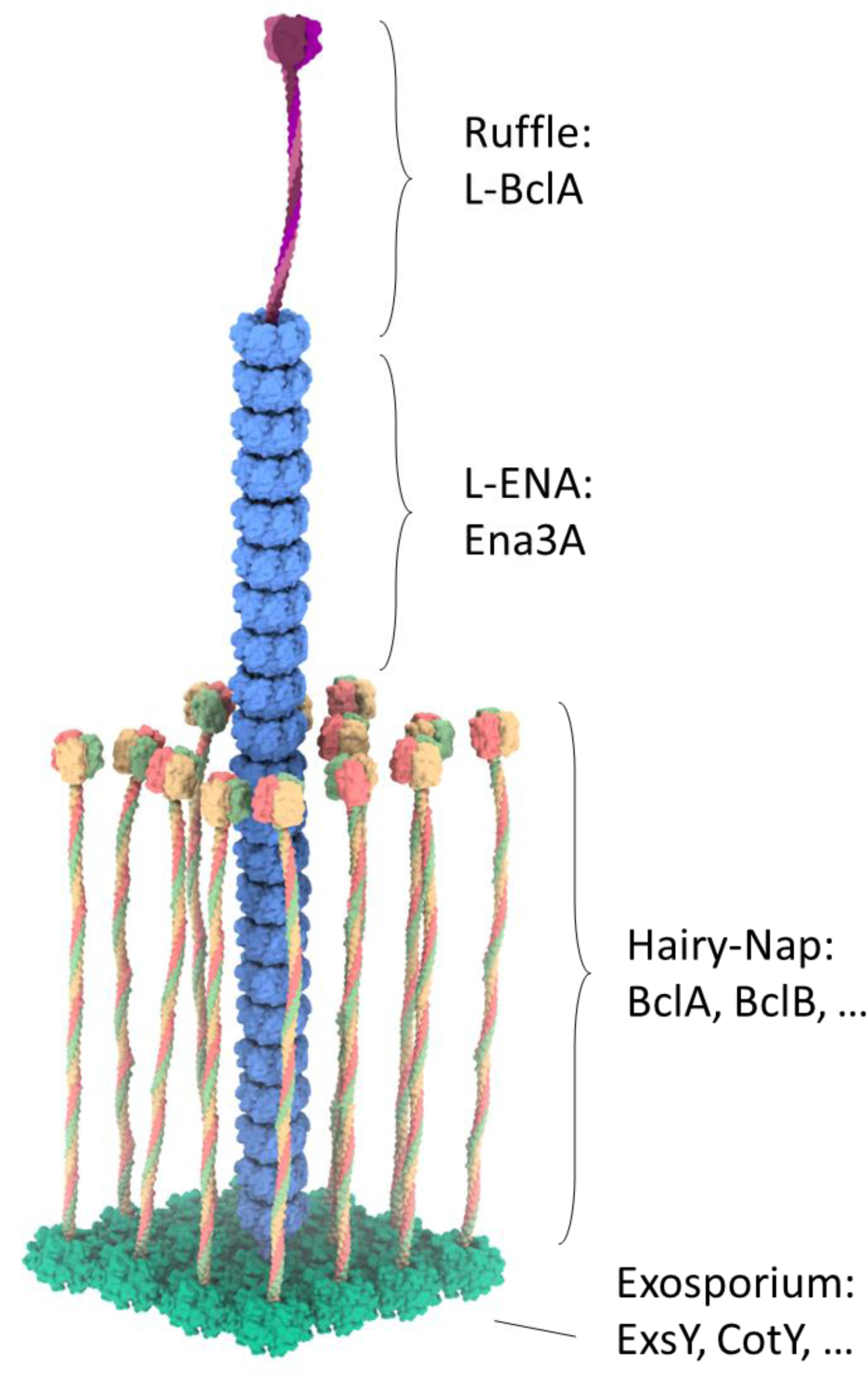
Mesoscale model of an L-ENA fiber projecting from the exosporium: L-ENA fibers are helical, protein ultrastructures composed of heptameric Ena3A rings that stack axially. At the tip that is distal from the spore, the L-ENA fiber terminates into a bacterial collagen-like ruffle protein, i.e., L-BclA. The L-ENA fiber protrudes through the hairy nap layer while being tethered to the exosporium via a dedicated anchoring protein CotZ. CotZ is expected to couple to or be integrated in the 2D crystal lattice of the exosporium. Note that for clarity, the L-ENA fibers are depicted shorter here.

## Discussion

In this contribution, we showed that the endospores of the food-poisoning outbreak strain *B. paranthracis* NVH 0075-95 are decorated with two types of filamentous protein structures, i.e., endospore appendages. Although prominently present on the endospore’s surface (*34*) and described since the 1970s for *Bacillus* (*10*), the biological role(s) of ENAs remain poorly understood (*18*). Following on the molecular identification of S-ENA (*17*), Jonsmoen *et al.* recently showed these fibers are involved in spore-spore aggregation (*19*). While the molecular underpinnings of the spore-spore coupling mechanism remains nuclear, we hypothesize that the S-ENA ruffle species is involved in establishing inter-spore interactions. Unfortunately, the genetic identity of the S-ENA ruffle remains unknown to date. Although structurally similar in appearance (i.e. a ∼2nm wide flexible fibrillum terminating in a knob-like domain), a difference in ruffle length and the *l-bclA* deletion phenotype (L-ENA ruffle: absent; S-ENA ruffle: present) show that the S-ENA ruffle protein is not encoded by *l-bcla*. This demonstrates that each ENA subtype is decorated by its own cognate ruffle protein, and hints at a functional diversification across the ENA subclasses. Knockout of the L-ENA subunit *ena3A* did not alter spore aggregation (*19*), reinforcing the notion that L-ENA fibers likely serve a specialized biological function that is distinct from spore clustering.

Can we infer a putative function for L-BclA? It is well established that Firmicutes endospores display bacterial collagen-like proteins (CLP) on their surface (*35–37*), but the biological function remains unclear for most of these proteins (*37*). The most well-known CLP within the *Bacillus* clade is the *<u>B</u>acillus* <u>c</u>ollagen-like protein of *<u>a</u>nthracis* (BclA) (*25*), the major glycoprotein (*38*) of the hairy nap (*39*) that is targeted to the exosporium via an N-terminal exosporium leader sequence (*31*). BclA has been shown to recruit complement factor H (CFH) to the spore surface (*40*), and its recruitment onto the exosporium facilitates C3 degradation, which in turn inhibits downstream complement activation, and ultimately promotes spore persistence. Next to its role in complement immune response evasion via CFH hijacking (*41*), BclA also facilitates spore entry into lung epithelial cells (*42*) via the receptor integrin α2β1 (*43*) and can also be used as an effective booster against inhalational anthrax in mouse models (*6*).

Similar to BclA, the CLP ruffle (i.e., L-BclA) found at the termini of the L-ENA fibers are found on the outermost layers of the endospore. Given the average length of the L-ENA fibers (±250nm), they extend well beyond the typical height of the hairy nap (±35nm) and therefore form one of the first points of contact of the spore to its environment. Speaking in broad terms, one could say that the spore has developed purpose-built ultra-stable display platforms (i.e., ENAs) to project specialized CLPs distally from the spore surface. In light of the fact that the parental NVH0075-95 strain is a cytotoxic foodborne outbreak isolate (*44*), combined with the observation that the L-ENA operon is found as mobile genetic elements and os exclusive to pathogenic *B. cereus* s.l. strains, and given the immuno-modulating role of BclA ‒one of L-BclA’s close structural homologues‒ it is tantalizing to speculate that L-ENA fibers play a role in spore virulence, be it either through e.g., direct binding to host epithelial surfaces, inhibition of phagocytosis, or by exerting additional control over the host immune system. Our structural understanding of the L-ENA / ruffle architecture combined with the genetic identification of all the pilus constituents, and the development of the three L-ENA gene cluster deletion strains (Δ*cotZ*, Δ*l-bclA*, Δ*ena3A*), has now paved the way for future experimental works to further help establish the biological role(s) of L-ENA fibers.

From a structural perspective, L-ENA fibers are remarkable products of macromolecular self-assembly, who’s assembly and surface display require a mere three genes: *cotZ*, *ena3A* and *l-bclA*. Each Ena3A subunit is cross-linked to its neighbor via 3 disulphide bridges, which totals to 21 covalent connections for each heptameric unit. Consequently, L-ENA fibers are able to withstand extreme physico-chemical stressors (heat, strong acids, chaotropes, detergent, and desiccation) and are among the most stable protein structures found in nature. In its biological context, however, a protein fiber will only be as strong as its weakest link. It is therefore expected that these high tensile strength ENAs require an anchoring mechanism of comparable durability. The phenotype of Δ*cotZ* spores (i.e. spores with detached L-ENA) combined with the AF2 modelling of the putative Ena3A-CotZ and CotZ-ExsY heteromers suggest that the anchoring complex is directly integrated into the exosporium 2D lattice. The exosporium of the *B. anthracis*/*cereus*/*thuringiensis* group is a flexible, disulphide cross-linked, protein 2D lattice that is predominantly composed of ExsY hexameric units (*45*). The AF2 model for the ExsY-CotZ dimer suggests that CotZ could be embedded within the ExsY lattice by mimicking the homotypic ExsY contacts, including the formation of an inter-molecular disulphide bridge. Similarly, the AF2 model for a CotZ-Ena3A dimer is reminiscent of the Ena3A contacts found in the L-ENA cryoEM model – this in turn also includes one of the intra-ring Ena3A disulphide bridges. Based on this, we hypothesize that CotZ acts as a bridging moiety that anchors the L-ENA pilus onto the spore surface by maintaining an unbroken chain of disulphide cross-links across the various contact points (i.e., ExsY → CotZ; Cotz → Ena3A). The experimental determination of the respective stoichiometries and the elucidation of the complex epitopes will be the subject of further study.

Astonishingly, L-ENA architecture marries an extreme mechanical resilience to a high degree of flexibility. That flexibility can be traced back to the segmented fiber ultrastructure wherein each ring section is coupled to the next via seven flexible N-terminal connectors. This enables rocking degrees of freedom between the neighboring ring segments without inducing steric clashes. The relative ease by which both S- and L-ENA fibers can be recombinantly produced, combined with their extraordinary material properties, makes them interesting starting points for the design of novel biobased materials. Despite the structural similarities (rmsd = 3.7Å) between Ena3A and Ena1B at the fold level, the L- and S-ENA pili have distinctly different topologies, i.e., ‘helical stairway’ versus ‘stacked rings’. By careful re-engineering of the dimer Ena-interface between the Ena subunits we anticipate that other ENA architectures could be (re)designed in service of their desired material properties.

## Materials and Methods

### Negative stain transmission electron microscopy

Negative stain TEM (nsTEM) imaging of bacterial spores and recEna3A filaments was done using formvar/carbon-coated copper grids (Electron Microscopy Sciences) with a 400-hole mesh. The grids were glow-discharged (ELMO; Agar Scientific) with 4mA plasma current for 45 seconds. 3 μl of a bacterial spore suspension or recEna3A protein solution was applied onto the glow-discharged grids and left to adsorb for 1 minute. The solution was dry blotted, followed by three washes with 15 μl Milli-Q. Next, 15 μl drops of 2% uranyl acetate were applied three times for 10 seconds, 2 seconds, and 1 minute respectively, with a blotting step in between each application. The excess uranyl acetate was then dry blotted with Whatman type 1 paper. All grids were screened with a 120 kV JEOL 1400 microscope equipped with LaB6 filament and TVIPS F416 CCD camera.

### Cryo-electron transmission microscopy

QUANTIFOIL® holey Cu 400 mesh grids with 2-µm holes and 1-µm spacing were glow discharged in vacuum using plasma current of 5 mA for 1 minute (ELMO; Agar Scientific). 3 µl of 0.6 mg/ml graphene oxide (GO) solution was applied onto the grid and incubated 1 min for absorption at room temperature. Excess GO was blotted using a Whatman grade 1 filter paper and left to dry under ambient conditions. For cryo-plunging, 3 µl of fiber suspension was applied on the GO-coated grids at 100% humidity and room temperature in a Gatan CP3 cryo-plunger. After 1 minute of absorption, the grid was machine-blotted with Whatman grade 2 filter paper for 3.5 seconds from both sides and plunge frozen into liquid ethane at −176°C. Grids were stored in liquid nitrogen until data collection. Two datasets were collected for *ex vivo* and recEna3A appendages with slight changes in the collection parameters. High-resolution cryoEM movies were recorded on a JEOL CRYO ARM 300 microscope equipped with omega energy filter and a K2 or K3 direct electron detector run in counting mode. For the *ex vivo* Ena, the microscope was equipped with a K2 summit detector and had the following settings: 300 keV, 100 mm aperture, 30 frames/image, 62.5 e−/Å^2^, total exposure 2.315-second exposure, and 0.82 Å/pixel. The recEna3A dataset was recorded using a K3 detector, at a pixel size of 0.782 Å/pix, and a total exposure of 64.66 e^−/^Å^2^ accrued over 61 frames/movie.

### Generation of deletion mutants and complementation constructs

Deletion mutants in *B. paranthracis* NVH0075-95 were generated following the marker less gene replacement method described previously (Janes and Stibitz, 2006; Pradhan et al., 2021). The gene replacement constructs contained the start and the stop codons of the respective genes to be deleted flanked by upstream and downstream homologous sequences. The upstream and downstream homologous sequences were 806 bp and 677 bp for *ena3A*, 727 bp and 750 bp for *l-bclA*, and 701 bp and 728 bp for *cotZ,* respectively. Synthetic constructs (Synbio Technologies LLC) were cut out from the pUC57-Amp vector by digesting with EcoRI, and the gene replacement DNA fragment were cloned into EcoRI digested pMAD-I-SceI shuttle plasmid (Lindbäck et al., 2012). The gene replacement constructs, i.e., pMAD-I-*Sce*I-Δ*ena3* and pMAD-I-*Sce*I-Δ*bcla*, and pMAD-I-*Sce*I-Δ*cotz* were then used to transform NVH0075-95 or the Δ*ena1ABC* triple mutant (Pradhan et al., 2021). Details of competent cell preparation, transformation, and screening are described (Zegeye and Aspholm, 2022; Pradhan et al., 2021).

For complementation experiments, DNA fragment containing a 300 bp region upstream of the start codon of the *l-bclA-ena3A operon* and open reading frames of the respective genes was ordered (Synbio Technologies LLC) or amplified by PCR and cloned in the low copy number plasmid pHT315 at EcoRI restriction site.

### Culturing and RNA extraction

*B. paranthracis* NVH0075-95 and *B. paranthracis* NVH0075-95 Δ*enaABC/*Δ*ena3A* quadruple mutant (control) were cultured in sporulation medium as previously described (Pradhan et al., 2021) with a slight modification. Here, we used nutrient broth (oxoid) in place of bacto medium (Difco). A sample of 20 ml culture was withdrawn every four hours from three independent cultures, centrifuged, and cell pellets frozen immediately at −80 °C until RNA extraction.

RNA was extracted using Purelink RNA minikit (Ambion, Life Technologies) according to the manufacturer’s protocol with a slight modification. Briefly, cell pellets were thawed on ice followed by addition of 200 µl of lysozyme solution (10 mM Tris-HCl (pH 8), 0.1 mM EDTA, 10 mg/ml lysozyme) and incubation for 5 minutes at RT. A volume of 1 µl of 10% SDS was added to the cell suspension and vortexed, after which 150 µl was transferred to new RNAse-free tubes. Following addition of 350 µl of lysis buffer, the cells were lysed by bead beating (FastPrep®-24, MP Biomedicals) for a total of two minutes at Speed 6, with 1 minute cooling on ice every 30 seconds. The lysates were centrifuged for 5 minutes (2600 xg), and the supernatants were transferred to new RNAse-free tubes. The remaining steps were carried out as described in the manufacturer’s protocol. RNA concentration was measured using Nanodrop 1000 (Thermo Scientific), and 1.5 µg RNA was treated with Turbo DNAase (Ambion) in 20 µl reaction volume following the manufacturer’s instruction.

### Quantitative real-time PCR (qRT-PCR) and PCR

Complementary DNA (cDNA) was synthesized from 500 ng DNAse-treated RNA using the QuantiTect reverse transcription kit (Qiagen) according to the manufacturer’s instructions. qRT-PCR was carried out using the PowerUp™ SYBR™ Green Master Mix kit (Applied Biosystems) and the AriaMx Real-Time PCR system (Agilent Technologies) following the manufacturers’ instructions. The expression of *cotZ*, *l-bclA*, *ena3A* and *rpoB* (housekeeping gene) was analyzed from cDNA of each of three independent cultures sampled after 4, 8, 12 and 16 hours after inoculation and the relative gene expression ratio was calculated according to the Pfaffl formula (Pfaffl, 2001). Primers were used at 300 nM concentration, and their efficiencies were calculated from the slope of the standard curve using the formula (10^-1/slope^-1) x 100, and were found to be between 95% and 101%. No template was added to the negative control samples. The Ct values obtained for the 4-hour samples were used as the calibrator sample in the data analysis, and *rpoB* served as an internal control gene. Each cDNA sample was analyzed in triplicate, and the experiment was repeated independently. The cycling conditions of the qRT-PCR were: 50 °C/2 minutes, 95°C/2 minutes; and 40 cycles of 95°C/15 seconds, 60°C/1 minute.

To determine if *cotZ*, *bclA*, and *ena3A* are co-expressed, standard PCR was conducted using the cDNAs as templates. Primers that span across the indicated genes (Table S4), and DreamTaq PCR master mix (Thermo Scientific™) were used. Purified genomic DNA (gDNA) from NVH0075-95 and cDNA from NVH0075-95 Δ*enaABC/*Δ*ena3A* (4 h and 16 h) were used as positive and negative controls, respectively.

### Sequencing strain *B. paranthracis* NVH 0075-95

To achieve a closed, updated, and high-quality genome of *B. paranthracis* NVH 0075-95, the strain was subjected to hybrid assembly of long- and short read sequences. Genomic DNA from NVH 0075-95 was prepared from bacteria grown over night on blood agar plates. All sequencing was performed by Novogene Co. (London, UK) using a combination of Pacific Biosciences (PacBio) RS II single-molecule real-time (SMRT) sequencing platform with a with a SMRTbell template library and an Illumina NovaSeq 6000 platform (2 x 150 bp) with an insert size of 300 bp.

A closed genome was achieved using the assembly pipeline Unicycler v. 0.4.8.0+galaxy3 (*46*) conducting a short-read-first hybrid assembly. Genome contiguity, completeness and correctness was assess using Quast v. 5.0.2+galaxy3 (*47*), Bandage 0.8.1+galaxy3 (*48*), BUSCO v. 5.3.2+galaxy0 (mode genome, gene predictor prodigal, lineage dataset bacillales_odb10) (*49*) and post-assembly correctios using Pilon ((*50*)) wrapped in the assembler Unicycler v. 0.4.8.0+galaxy3. Species validation was done on the Type Strain Genome Server (TYGS) (*51, 52*). Plasmids and contigs were predicted with PlasFlow v. 1.1.0 (threshold >0.6, remaining default settings) (*53*), and the genome was annotated using the online NCBI Procaryotic Genome Annotation Pipeline (PGAP) (*54–56*). Data is available at NCBI under accession number GCA_027945115.1.

### Distribution of *L-ena*

Cblaster ((*57*)) a python toolkit for detecting collocated genes, was run remotely through the web application CAGECAT v.1.0 (settings: -max gap 200 bp, n of unique query seq=3, min coverage 50%, min identity 30%, maximum e-value 0.01) on the NCBI RefSeq non-redundant protein database, using amino acid sequence of WP_017562367.1 (Ena3A), WP_048548723.1 (BclA), WP_048548726.1 (CotZ) and WP_058548727.1 (Tn3 family transposase) as query. The genomic region spanning WP_017562367.1, WP_048548723.1, WP_048548726.1 is hereafter referred to as “the gene cluster”. Furthermore, to investigate the proportion of organisms with the gene cluster within the *B. cereus s.l.* group, publicly available genomes yielding hits in the cblaster search (Appendix Table) were downloaded from NCBI RefSeq database (n = 656, NCB (https://www.ncbi.nlm.nih.gov/refseq/; Table EV1) and appended to a representative database of assemblies of *B. ceresu s.l.* group (Supplementary Table 1). In addition, 136 *B. subtilis* were included for comparison. Assemblies were quality checked using QUAST, and only genomes of correct size (∼4.9 – 6.2 Mb) and a GC content of ∼35% were included in the downstream analysis. Pairwise tBLASTn searches were performed (BLAST+ v. 2.11.0 ((*58*)), e-value 1e-10, max_hspr 1, default settings) to search for homo- and orthologs of the following query protein sequences from strain NVH 0075-95: WP_017562367.1, WP_048548723.1 and WP_048548726.1. Proteins were considered as orthologs or homologs when they matched the query protein with high coverage (> 70%) and moderate sequence identity (> 30%), and when the whole gene cluster was present in the genome with corresponding synteny as the NVH 0075/95 strain. Genomic location (chromosome or plasmid) was inspected manually for complete and closed genomes. Mashtree v. 0.57 ((*59*)) was used to infer whole genome clustering for the *B. cereus s.l.* group, using the accurate option (--min-depth 0). Clustering and metadata (hits for query proteins) were visualized in Microreact web browser ((*60*)).

---

**Supporting Figure 1:**
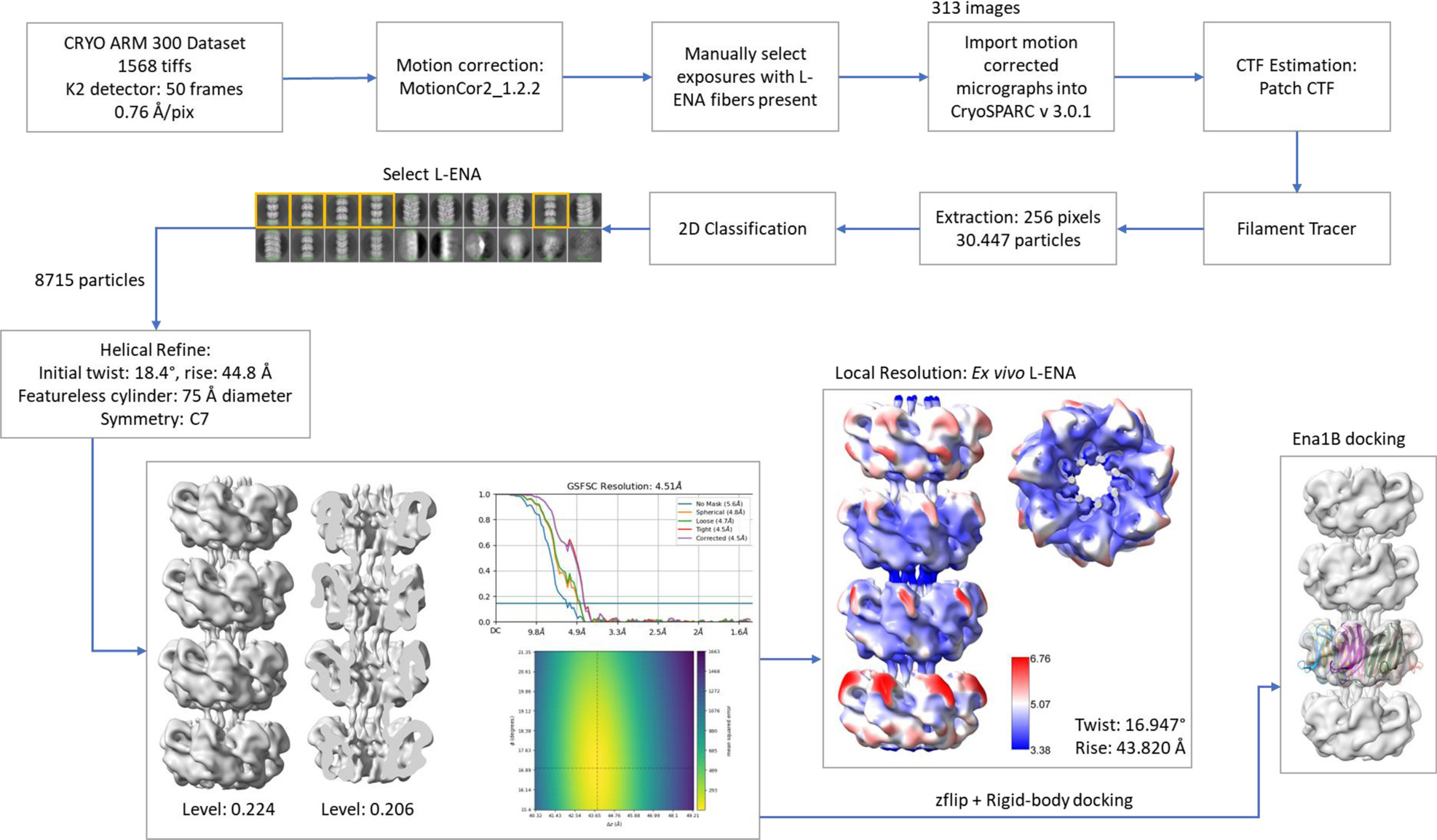
CryoEM workflow for the *ex vivo* L-ENA fibers

**Supporting Figure 2:**
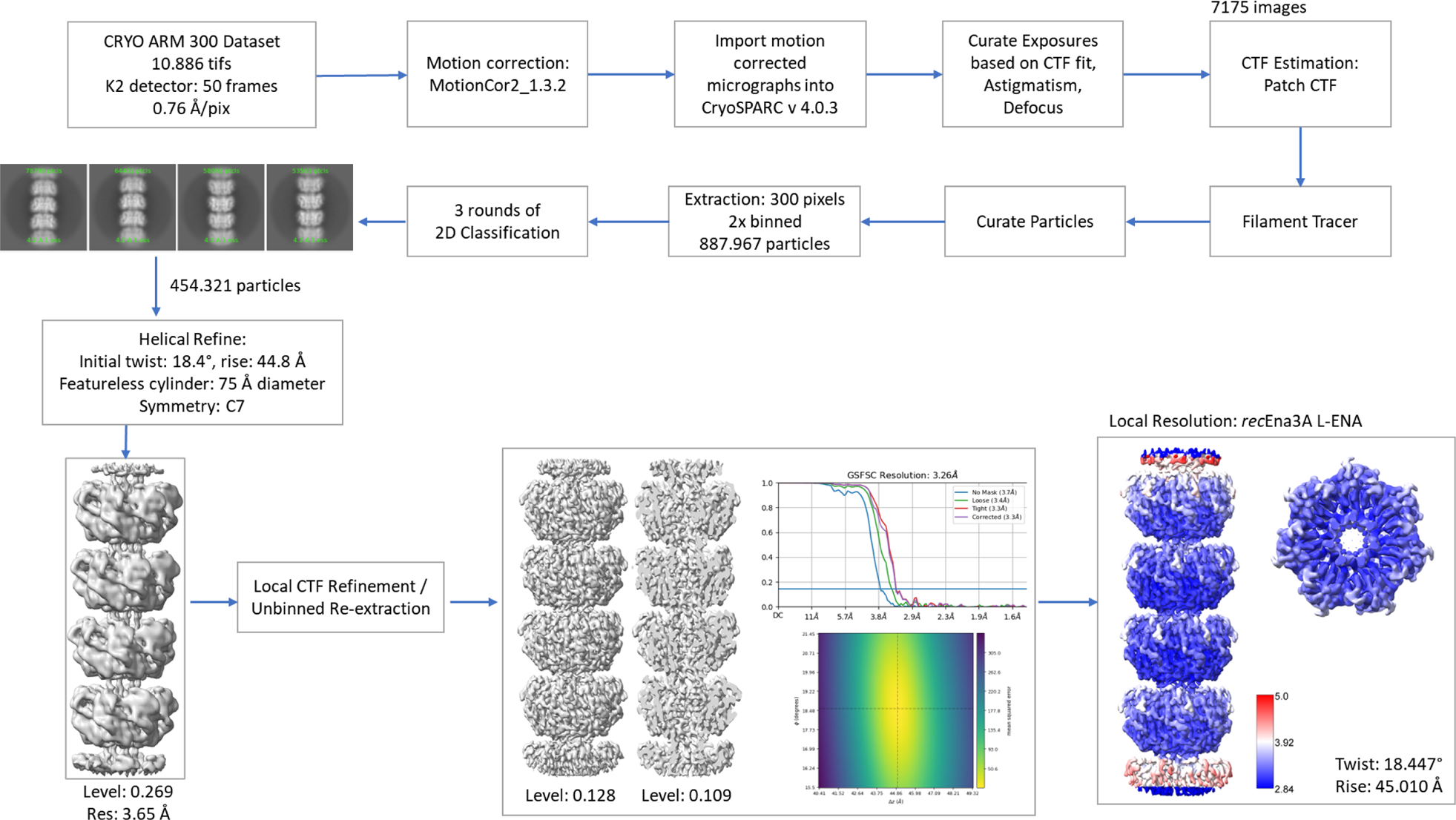
CryoEM workflow for the *rec*Ena3A L-ENA fibers

**Supporting Figure 3:**
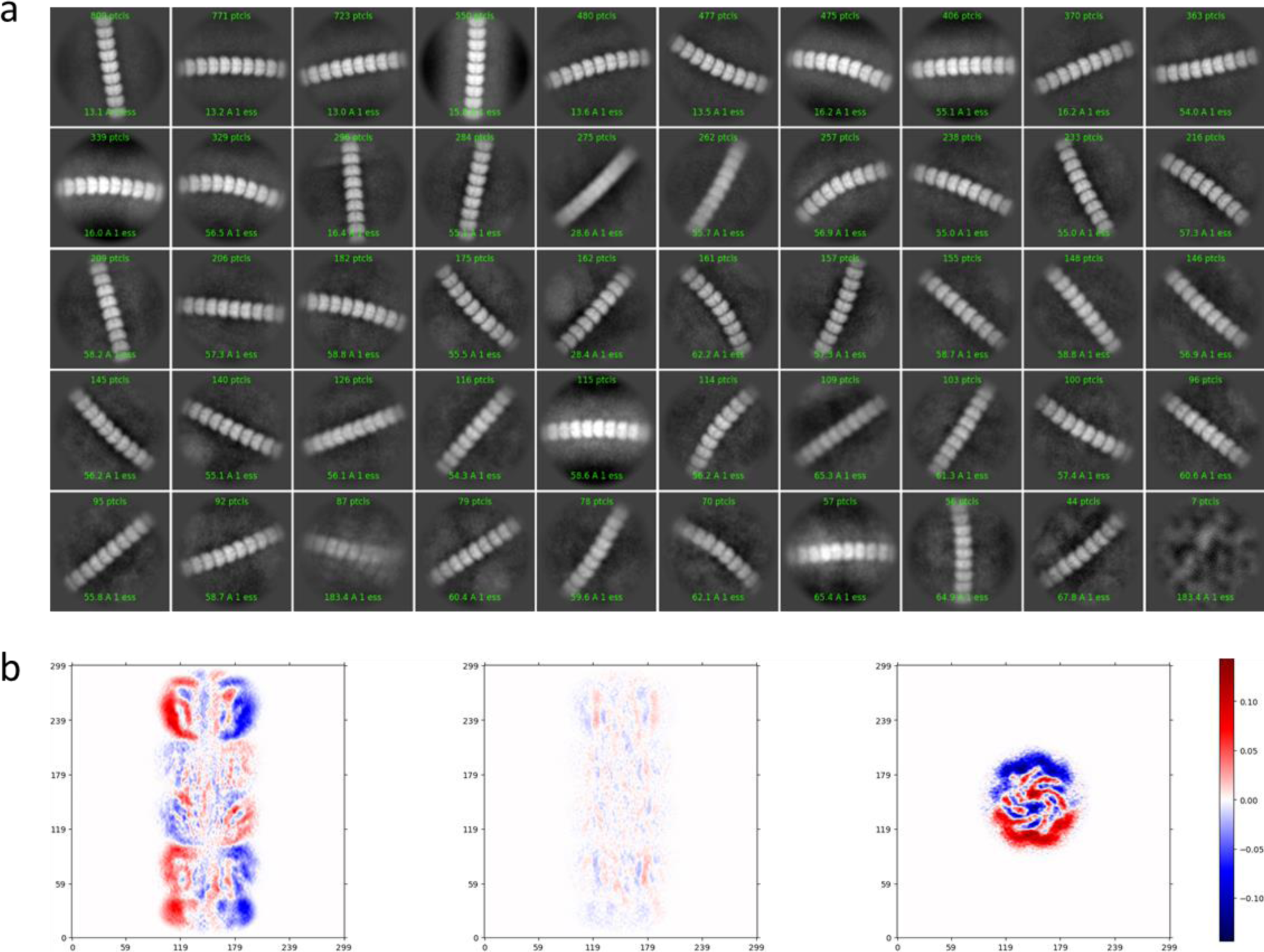
Flexibility of *rec*Ena3A L-ENA fibers: (a) CryoSPARC 2D class averages of L-ENA using a particle box size of 600×600pixels (0.76Å/pixel) after manual picking of high curvature L-ENA segments, (b) Cryosparc output for the mode 1 of the 3D variability analysis job on L-ENA (see Supporting Movie 1 for a representation of the corresponding volume series).

**Supporting Figure 4:**
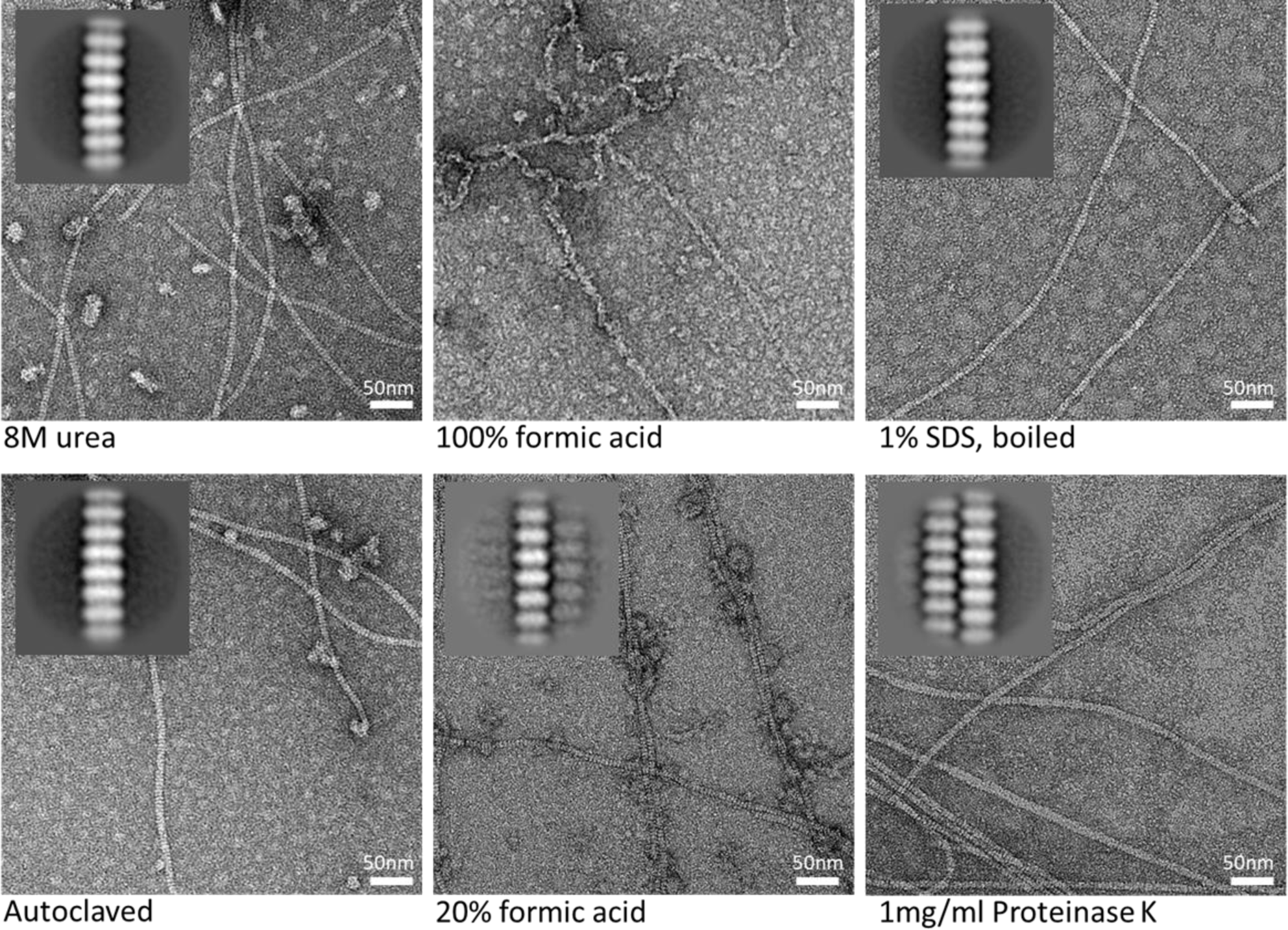
Stability assay for recEna3A fibers: recombinant Ena3A fibers were incubated for 1h in 8M urea, 20% or 100% (v/v) formic acid, or boiled in 1% (w/v) SDS, autoclaved (121C° for 18min) in miliQ, or incubated in 1 mg.mL^-1^ proteinase K in 33.3mM Hepes pH 7.5, 1mM CaCl_2_ for 24h.

**Supporting Figure 5:**
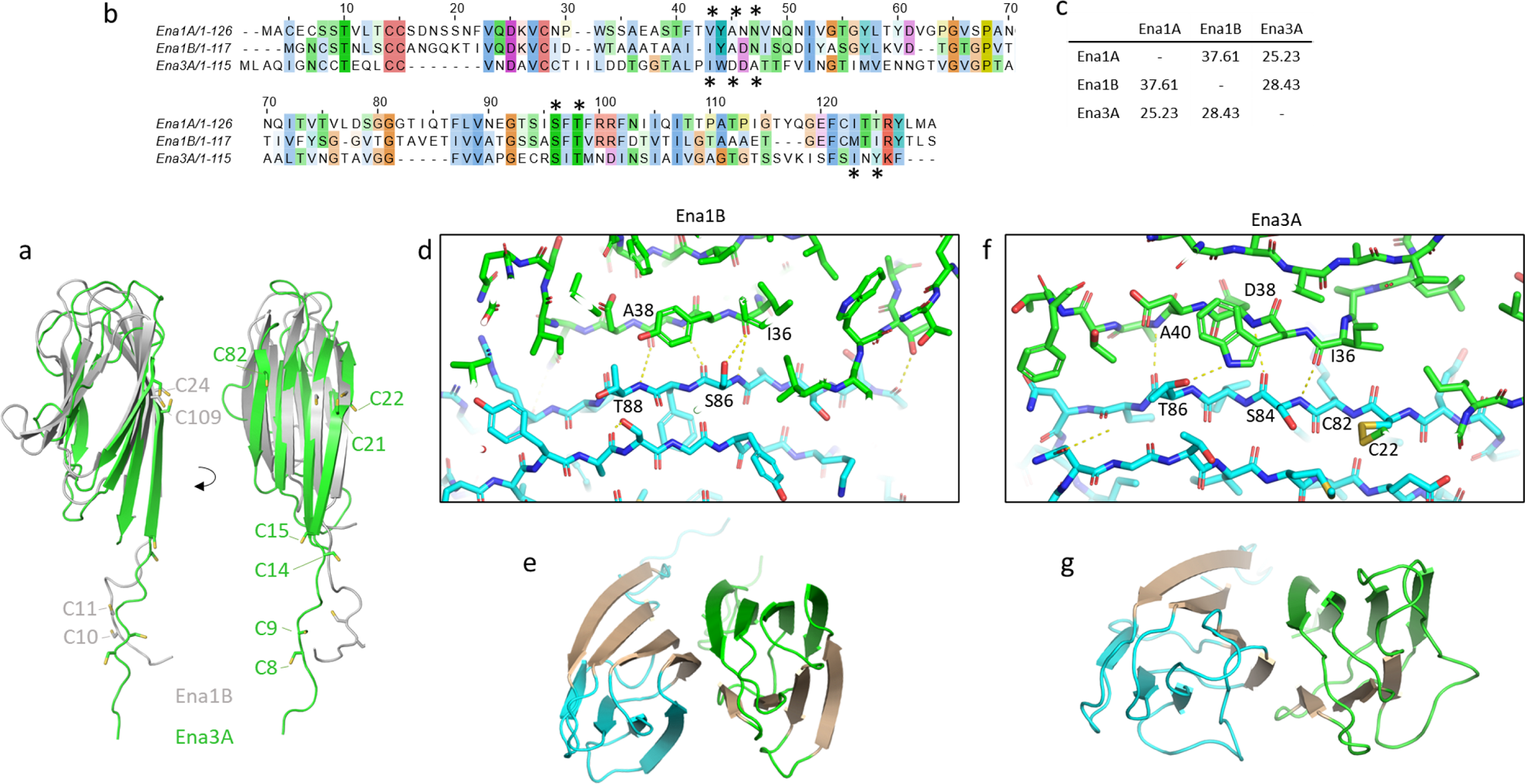
Multiple sequence alignment of the S-ENA subunits (Ena1A and Ena1B) and the L-ENA subunit (Ena3A). Residues involved in lateral contacts are highlighted with a star (above: S-ENA contact; below: L-ENA contacts), (b) percent identity matrix for Ena1A, Ena1B and Ena3A, (c) and (e) comparison between the β-sheet augmentation contacts at the Ena1B dimer interface (PDB: 7A02), and (d) and (f) the β-sheet augmentation at the Ena3A dimer interface. Inter-molecular hydrogen bonds (determined in PyMol 2.5.2 using “Find Polar between Chains”) shown in yellow, dashed lines.

**Supporting Figure 6:**
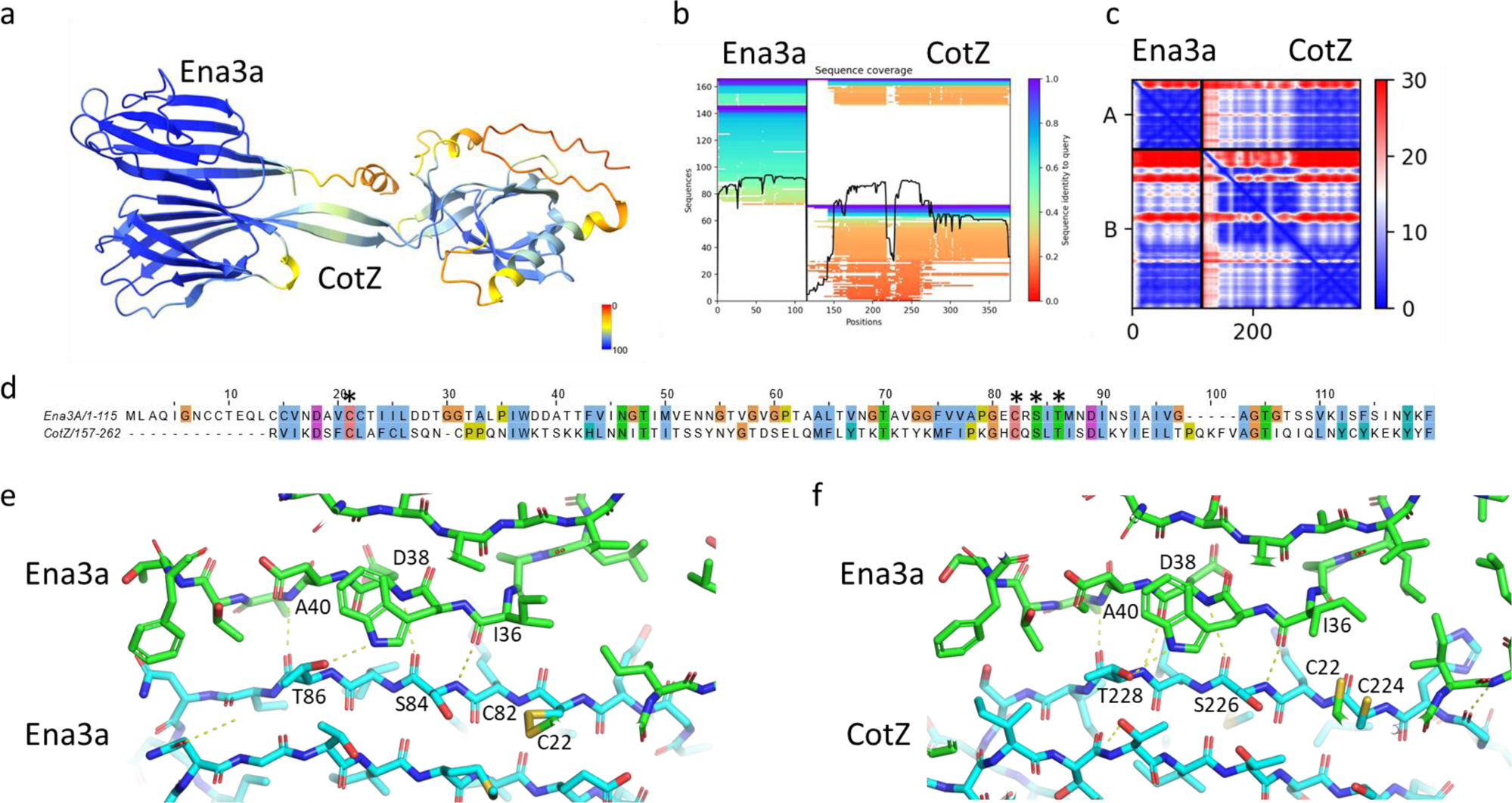
Alphafold2 prediction of the CotZ-Ena3A dimeric complex: (a) Alphafold-multimer 1.2 Ena3A-CotZ dimer (pLDDT=82.6; ptmscore=0.73) colour coded according to pLDDT-score, (b) Sequence coverage of the corresponding Ena3A-CotZ multiple sequence alignment, (c) Predicted aligned error map of the predicted Ena3A-CotZ dimer, (d) Pairwise sequence alignment (17.4% sequence identity) of Ena3A and the C-terminal Ena-core domain of CotZ (157-262). Residues involved in lateral Ena3A-Ena3A contacts (highlighted with a star) are conserved in CotZ, (e) β-sheet augmentation at the Ena3A dimer interface (determined via cryoEM), (f) Predicted β-sheet augmentation at the Ena3A-CotZ heterodimer interface (predicted using AF2 multimer). Inter-molecular hydrogen bonds (determined in PyMol 2.5.2 using “Find Polar between Chains”) shown in yellow, dashed lines.

**Supporting Figure 7:**
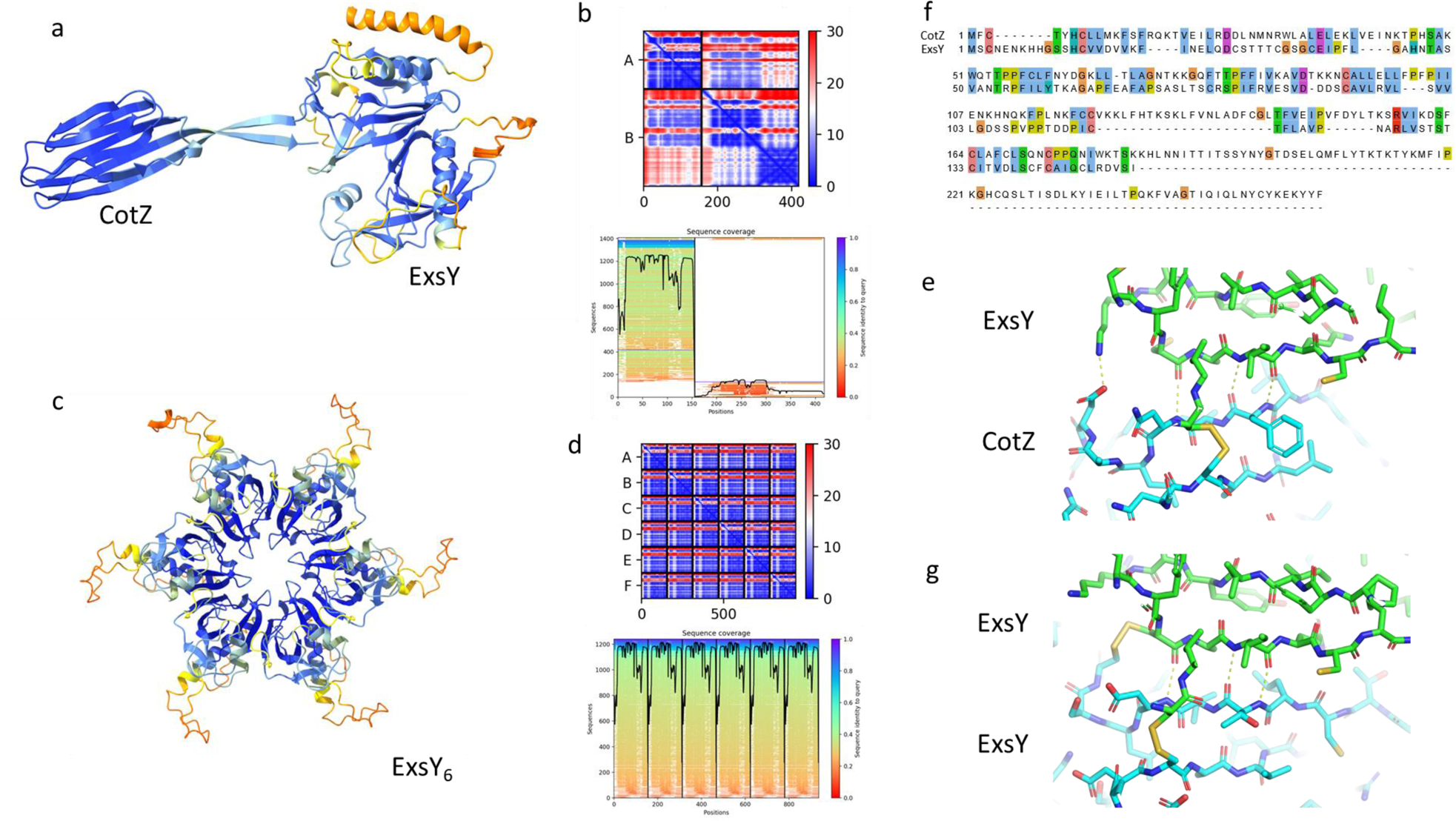
(a) Alphafold-multimer 1.2 CotZ-ExsY dimer (pLDDT=80.2; ptmscore=0.71) colour coded according to pLDDT-score, (b) Predicted aligned error map and sequence coverage of the multiple sequence alignment, (c) Alphafold-multimer 1.2 ExsY hexamer (pLDDT=81.3; ptmscore=0.86) colour coded according to pLDDT-score, (d) Predicted aligned error map and sequence coverage of the multiple sequence alignment, (e) Pairwise-sequence alignment between CotZ and ExsY (30.8% sequence identity), comparison of the AF2, (f) and (g) comparison of the putative ExsY/CotZ and ExsY/ExsY dimeric interfaces. Inter-molecular hydrogen bonds (determined in PyMol 2.5.2 using “Find Polar between Chains”) shown in yellow, dashed lines.

**Supporting Figure 8:**
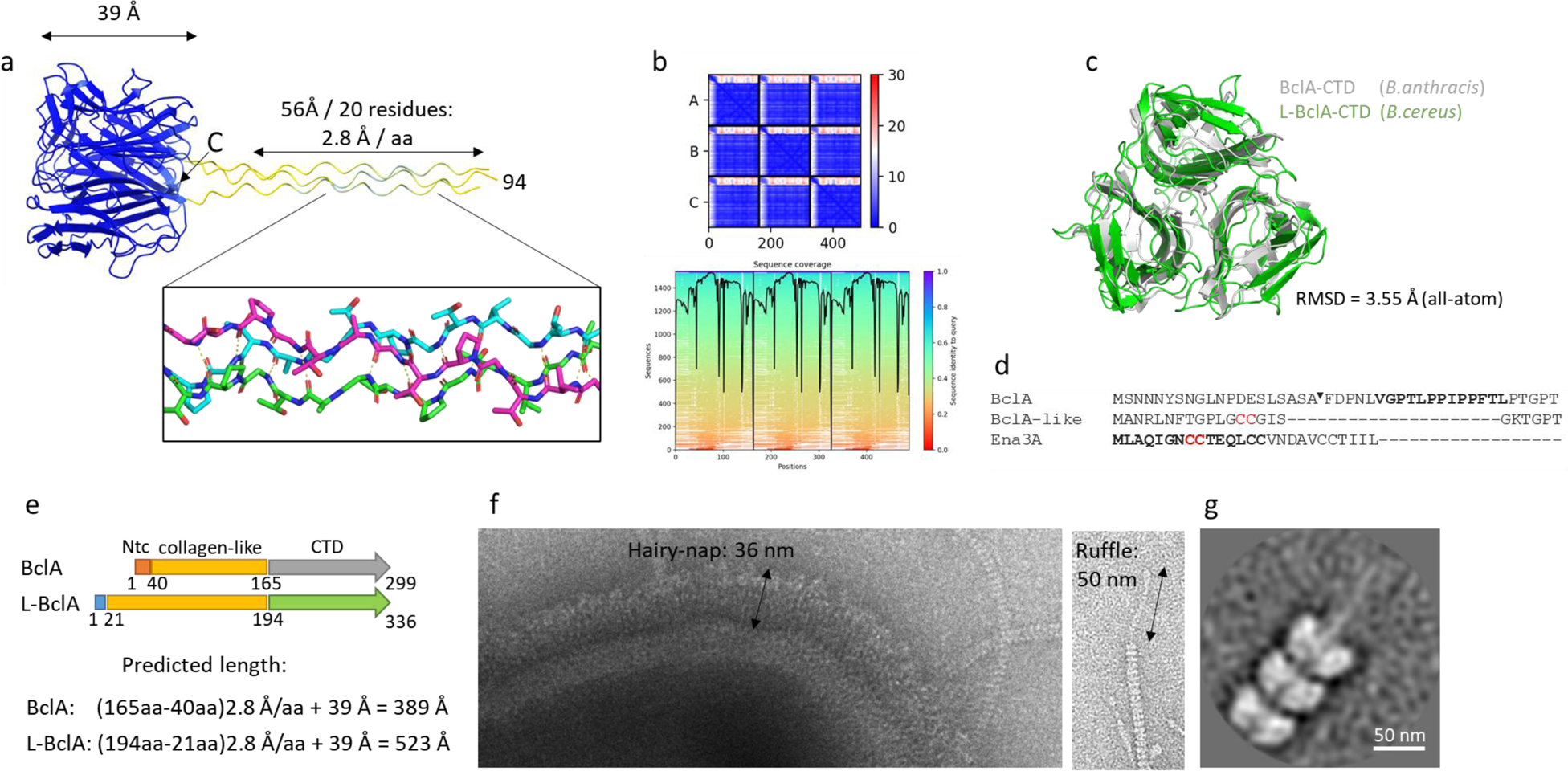
Structural analysis of L-BclA: (a) Alphafold-multimer v 1.2 prediction of an L-BclA_94-267_ trimer (pLDDT=94, ptmscore=0.9) colour coded according to pLDDT-score (N-terminus omitted); collagen-like stalk region shown in stick representation with putative H-bonds shown in dashed lines, (b) Corresponding predicted aligned error map and sequence coverage of the multiple sequence alignment of the AF2 L-BclA_94-267_ trimer, (c) superposition (all-atom RMSD = 3.55Å; sequence identity = 22.1%) of the AF2 L-BclA_94-267_ trimer with the crystal structure of BclA-CTD of the hairy nap layer of *B.anthracis* (PDB: 1WCK), (d) N-terminal sequence of BclA, L-BclA and Ena3A: letters in bold for BclA correspond to the exosporium leader sequence, and the arrow marker indicates the proteolytic cleavage site. L-BclA has no identifiable exosporium leader sequence, nor any notable sequence homology to the N-terminal connector (shown in bold) of Ena3A apart from a single CC-motif, (e) Domain organization of BclA (Q81JD7) and L-BclA and the corresponding theoretical lengths in fully extended state, (f) Comparison of the thickness of the hairy nap layer to the length of the L-ENA ruffles, (g) 2D class average of L-ENA fiber termini.

**Supporting Figure 9:**
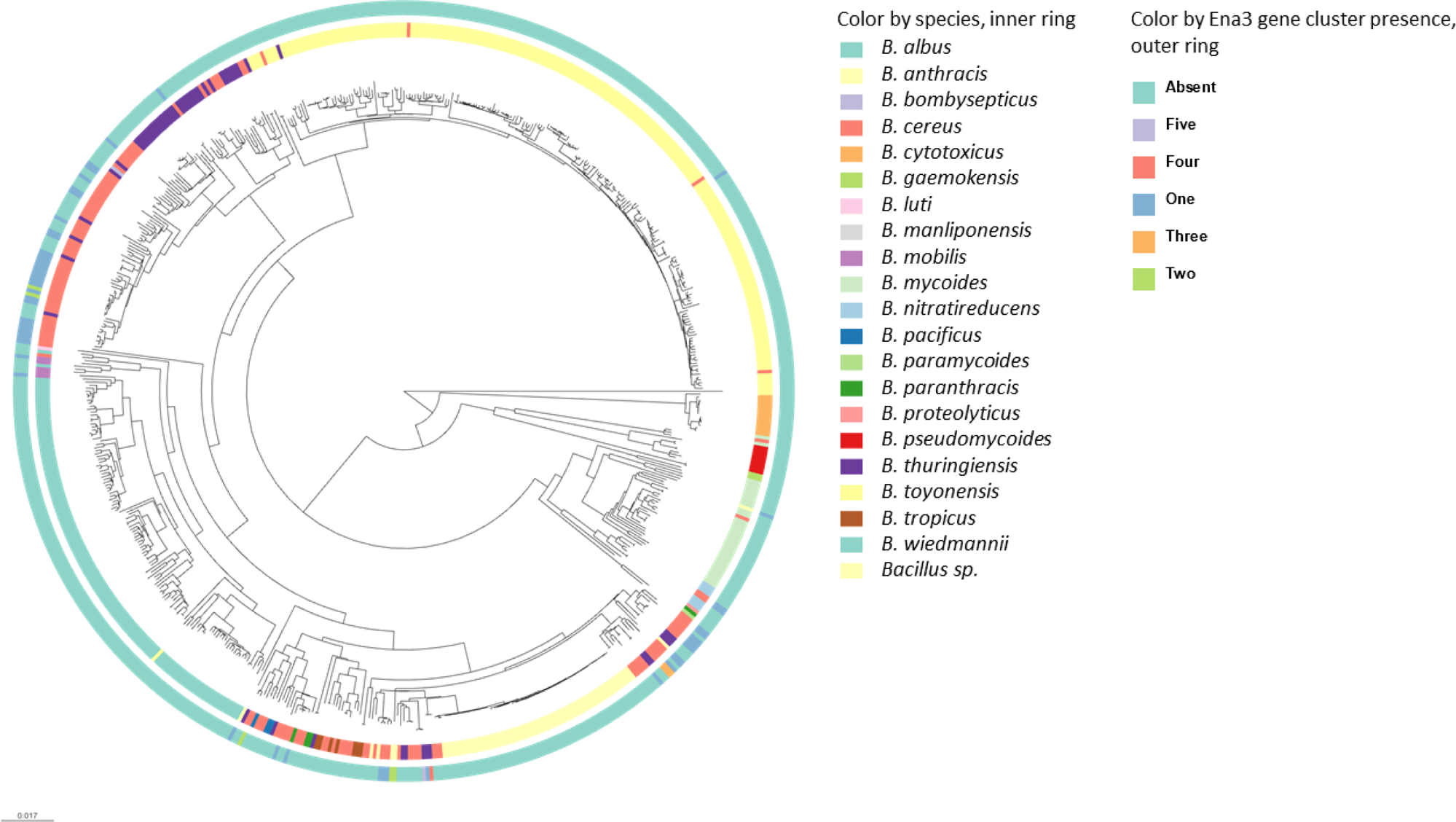
Phylogenetic analysis of L-ENA occurrence. Clustering of 656 *B. cereus s.l.* genomes (*B. subtilis* is excluded from tree). Rings are colored according to designated species (inner ring) and presence and copy-number of *ena3A* gene cluster (outer ring). Tree is made using Mashtree, vizualised using Microreact, and available at https://microreact.org/project/uzm4JFrrsCPZeRnMpRqvvf-supplementary-figure-9-ena3-paper.

**Table S1.**
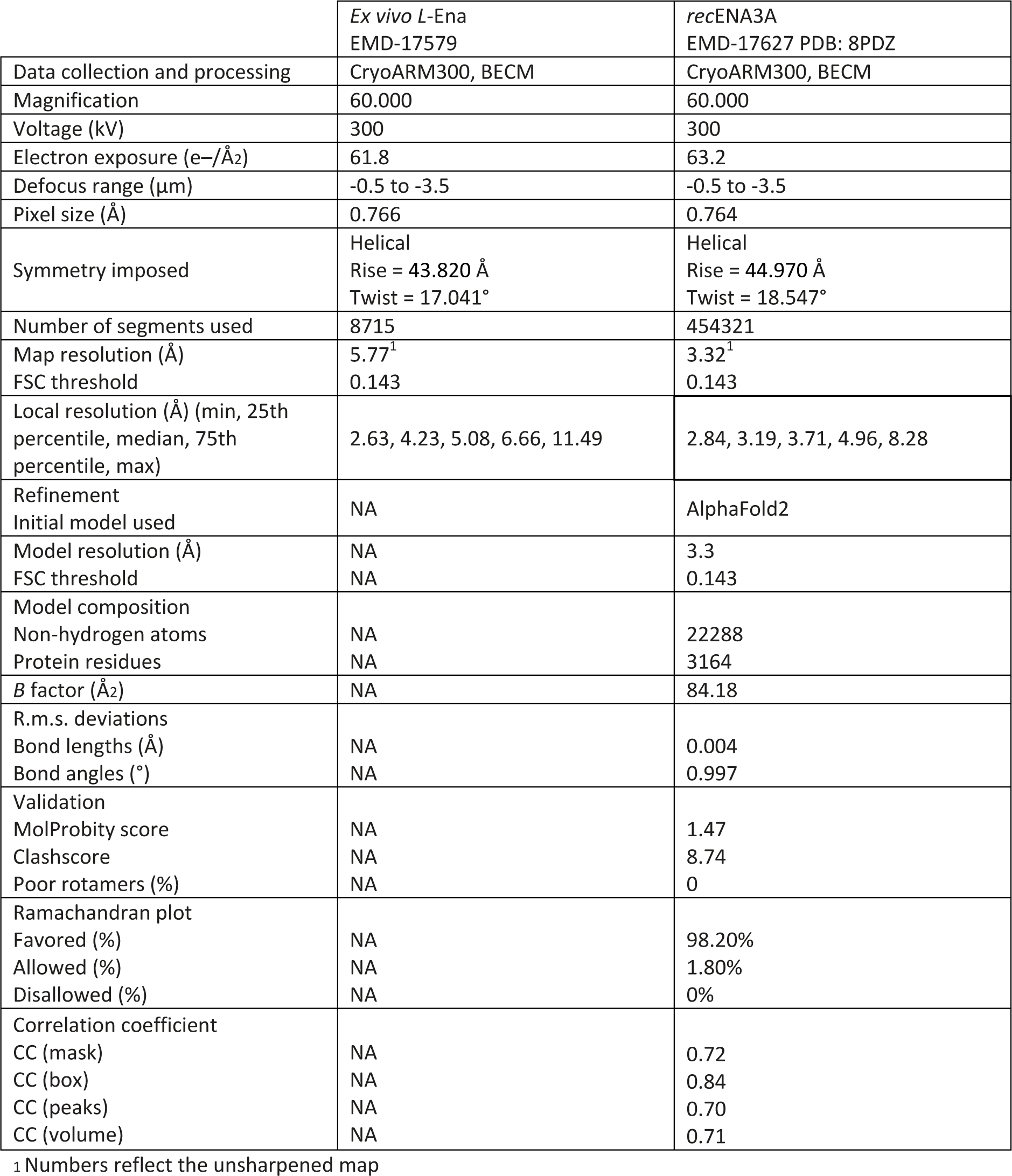
CryoEM model and data statistics.

|  | *Ex vivo* L-Ena<br>EMD-17579 | *rec*ENA3A<br>EMD-17627 PDB: 8PDZ |
| --- | --- | --- |
| Data collection and processing | CryoARM300, BECM | CryoARM300, BECM |
| Magnification | 60.000 | 60.000 |
| Voltage (kV) | 300 | 300 |
| Electron exposure (e-/Å<sup>2</sup>) | 61.8 | 63.2 |
| Defocus range (µm) | -0.5 to -3.5 | -0.5 to -3.5 |
| Pixel size (Å) | 0.766 | 0.764 |
| Symmetry imposed | Helical<br>Rise = 43.820 Å<br>Twist = 17.041° | Helical<br>Rise = 44.970 Å<br>Twist = 18.547° |
| Number of segments used | 8715 | 454321 |
| Map resolution (Å) | 5.77<sup>1</sup> | 3.32<sup>1</sup> |
| FSC threshold | 0.143 | 0.143 |
| Local resolution (Å) (min, 25th percentile, median, 75th percentile, max) | 2.63, 4.23, 5.08, 6.66, 11.49 | 2.84, 3.19, 3.71, 4.96, 8.28 |
| Refinement |  |  |
| Initial model used | NA | AlphaFold2 |
| Model resolution (Å) | NA | 3.3 |
| FSC threshold | NA | 0.143 |
| Model composition |  |  |
| Non-hydrogen atoms | NA | 22288 |
| Protein residues | NA | 3164 |
| B factor (Å<sup>2</sup>) | NA | 84.18 |
| R.m.s. deviations |  |  |
| Bond lengths (Å) | NA | 0.004 |
| Bond angles (°) | NA | 0.997 |
| Validation |  |  |
| MolProbity score | NA | 1.47 |
| Clashscore | NA | 8.74 |
| Poor rotamers (%) | NA | 0 |
| Ramachandran plot |  |  |
| Favored (%) | NA | 98.20% |
| Allowed (%) | NA | 1.80% |
| Disallowed (%) | NA | 0% |
| Correlation coefficient |  |  |
| CC (mask) | NA | 0.72 |
| CC (box) | NA | 0.84 |
| CC (peaks) | NA | 0.70 |
| CC (volume) | NA | 0.71 |

<sup>1</sup> Numbers reflect the unsharpened map

**Table S2.** Plasmid constructs.

| Name | Details | Reference |
| --- | --- | --- |
| *pMAD-I-SceI* | Shuttle vector for making deletion mutant, carries I-SceI restriction site | (Lindbäck et al., 2012) |
| *pMAD-I-SceI Δ*ena3A** | *pMAD-I-SceI* carrying homology sequences flanking *ena3A* | this study |
| *pMAD-I-SceI Δ*l-bclA** | *pMAD-I-SceI* carrying homology sequences flanking *l-bclA* | this study |
| *pMAD-I-SceI Δ*cotZ** | *pMAD-I-SceI* carrying homology sequences flanking *cotZ* | this study |
| *pBKJ233* | Expresses I-SceI enzyme (promotes second cross over) | (Janes and Stibitz, 2006) |
| *pHT315* | Low-copy number plasmid for complementation | (Arantes and Lereclus, 1991) |
| *pena3A* | *pHT315::ena3A* (complementation of *B. cereus* Δ*ena3A*) | this study |
| *pl-bclA* | *pHT315::l-bclA* (complementation of *B. cereus* Δ*l-bclA*) | this study |
| *pcotZ* | *pHT315::cotZ* (complementation of *B. cereus* Δ*cotZ*) | this study |

**Table S3.** Overview of gene knockout mutants/complementation constructs.

| **Name** | **Genetic background** | **Reference** |
| --- | --- | --- |
| Wild type | NVH0075-95 | (Pradhan et al., 2021) |
| $\Delta cotZ$ | NVH0075-95 | this study |
| $\Delta l-bclA$ | NVH0075-95 | this study |
| $\Delta ena3A$ | NVH0075-95 | this study |
| $\Delta ena1ABC$ | | (Pradhan et al., 2021) |
| $\Delta ena1ABC/\Delta ena3A$ | NVH0075-95 $\Delta ena1ABC$ | this study |
| $\Delta cotZ::pcotZ$ | NVH0075-95 $\Delta cotZ$ | this study |
| $\Delta bclA::pl-bclA$ | NVH0075-95 $\Delta l-bclA$ | this study |
| $\Delta ena3A::pena3A$ | NVH0075-95 $\Delta ena3A$ | this study |
| $\Delta enaABC/\Delta ena3A::pena3A$ | NVH0075-95 $\Delta ena1ABC/\Delta ena3A$ | this study |

**Table S4.**
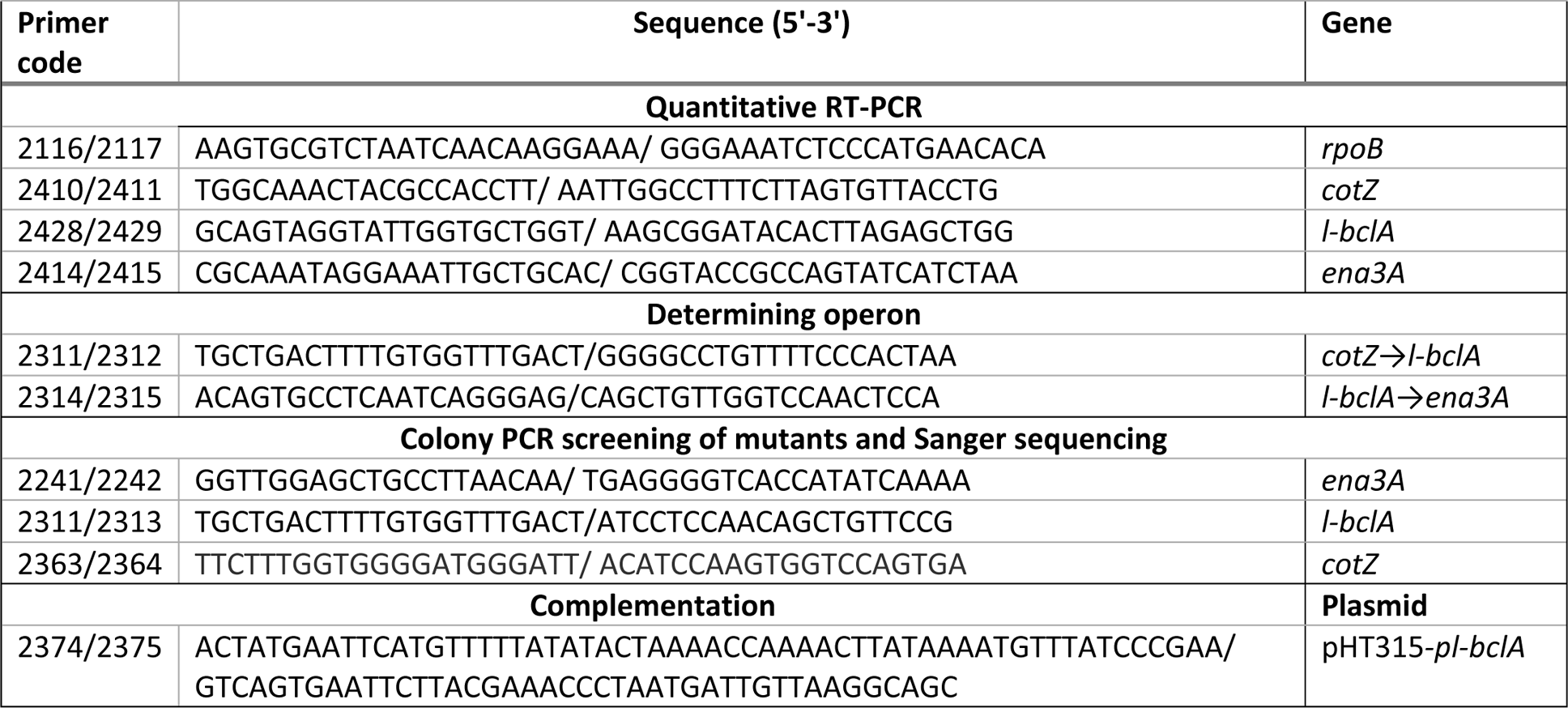
List of primers.

## Author contributions

MS, MA and HR designed the project. MS performed cryogenic freezing, nsTEM and cryo-EM imaging and data processing with assistance from MF. EDZ conducted the gene expression studies, designed and generated the knockout strains and complementation constructs, and prepared the spores for analysis. MS wrote the manuscript with contributions from all authors.

## Competing Interests Statement

The authors declare no competing interests.

## Data Availability Statement

The *ex vivo* L-ENA and the recombinant Ena3A cryo-EM maps were deposited to EMDB under entry IDs EMD-17579 and EMD-17627, respectively. The atomic model for recEna3A was deposited to the PDB under ID 8PDZ.

## Acknowledgements

We thank Dirk Reiter at the VIB-VUB Facility for Bio Electron Cryogenic Microscopy (BECM) and Yohannes Beyene Mekonnen at NMBU for technical assistance. This work was funded by VIB, NMBU, EOS Excellence in Research Program by FWO through grant G0G0818N to HR and G043021N to MS. MA recognizes the Grant from the Norwegian research council (NFR): 335029 - FORSKER22.

